# Full-length RNA transcript sequencing traces brain isoform diversity in house mouse natural populations

**DOI:** 10.1101/2024.01.03.573993

**Authors:** Wenyu Zhang, Anja Guenther, Yuanxiao Gao, Kristian Ullrich, Bruno Huettel, Aftab Ahmad, Lei Duan, Kaizong Wei, Diethard Tautz

## Abstract

The ability to generate multiple RNA transcript isoforms from the same gene is a general phenomenon in eukaryotes. However, the complexity and diversity of alternative isoforms in natural populations remain largely unexplored. Using a newly developed full-length transcripts enrichment protocol with 5’ CAP selection, we sequenced full-length RNA transcripts of 48 individuals from outbred populations and subspecies of *Mus musculus*, and from the closely related sister species *Mus spretus* and *Mus spicilegus* as outgroups. The dataset represents the most extensive full-length high-quality isoform catalog at the population level to date. In total, we reliably identified 117,728 distinct isoforms, of which only 51% were previously annotated. We show that the population-specific distribution pattern of isoforms is phylogenetically informative and reflects the segregating SNP diversity between the populations. We find that ancient housekeeping genes are a major source of the overall isoform diversity, and that the generation of alternative first exons plays a major role in generating new isoforms. Given that our data allow us to distinguish between population-specific isoforms and isoforms that are conserved across multiple populations, it is possible to refine the annotation of the reference mouse genome to a set of about 40,000 isoforms that should be most relevant for comparative functional analysis across species.

## Introduction

The ability to generate multiple RNA isoforms from the same gene increases vastly the complexity of the transcriptome and proteome in eukaryotes (Keren et al. 2010; Nilsen and Graveley 2010). Alternative isoforms with different coding sequences can be generated through multiple types of alternative splicing events (Naftaly et al. 2021), including skipped exon (SE), retained intron (RI), mutually exclusive exons (MX), alternative 5’ (A5’), and 3’ (A3’) splice sites. Isoform diversity can also be expanded via the inclusion of alternative transcription start sites (TSSs) or transcript end sites (TESs), resulting in differences in the untranslated regions (UTRs) (Landry et al. 2003; Tian and Manley 2017).

By analyzing high-throughput EST and short-read RNA-Seq data across species, several comparative studies on the complexity of isoform landscape have been performed in plants (Ling et al. 2019), insects (Malko et al. 2006), vertebrates (Mudge et al. 2011; Barbosa-Morais et al. 2012), birds (Rogers et al. 2021), mice (Harr and Turner 2010), and primates (Lin et al. 2010). These initial comparative studies have provided important insights into the enormous diversity of alternative isoforms, but only at the local level of splicing junction events due to the limitation of the short-read sequencing technology.

The recent development of long-read single-molecule sequencing technologies has enabled the capture of the full-length isoform diversity (Byrne et al. 2019), further facilitating the comparative analysis of alternative isoforms in different species. Leung SK et al. (Leung et al. 2021) used long-read isoform sequencing to generate full-length transcript sequences in the human and mouse cortex. They showed a similar global pattern of isoform diversity between humans and mice but with striking differences in some genes between these two species. Another recent study from Ferrandez-Peral L (Ferrandez-Peral et al. 2022) generated an extensive full-length isoform catalog in primates and found that a high isoform diversity in immune genes correlates with signals of positive selection in the coding regions of the genes. These between-species comparison studies have provided insights into the complexity of alternative isoforms at a deep evolutionary time scale. However, the diversity and pattern of the alternative isoform landscape within natural populations remain largely unexplored (Verta and Jacobs 2022).

Owing to its well-defined evolutionary history (Guenet and Bonhomme 2003; Phifer-Rixey and Nachman 2015), the house mouse (*Mus musculus*) is a particularly suitable model system for studying the dynamics of polymorphisms and recently originated genetic elements in natural populations. Currently, three major lineages of the house mouse, which diverged roughly half a million years ago, are distinguished as subspecies (Harr et al. 2016): the Western European house mouse *Mus musculus domesticus*, the Eastern European house mouse *Mus musculus musculus*, and the Southeast Asian house mouse *Mus musculus castaneus.* With a divergence time of fewer than 2 million years, closely related outgroup species (*e.g.*, *Mus spretus*) are also available in this model system (Harr et al. 2016). We have previously used the house mouse system to study the evolutionary pattern of single nucleotide variants (Staubach et al. 2012), gene copy number variants (Pezer et al. 2015), and gene retrocopy variants (Zhang et al. 2021; Zhang and Tautz 2022). This made the house mouse an ideal system to directly trace the origin and pattern of alternative isoforms in natural populations.

Here, we use mice brains as the source tissue for the transcriptome analysis. The brain harbors the largest diversity of cell types with an overall transcript diversity only comparable to testis (Bekpen et al. 2018), but with many more of these transcripts being likely to be functional in the brain compared to the testis, where there is a lot of expression due to a transcriptionally permissive chromatin environment, especially in late spermatogenic cell types (Murat et al. 2023). Previous studies have made use of long-read sequencing to discover the isoform diversities in cell types and developmental stages of the laboratory mouse inbred strain C57Bl6 (Gupta et al. 2018; Lebrigand et al. 2020; Joglekar et al. 2021; Ding et al. 2022). However, the brain isoform diversity in wild mice, especially across natural populations, has so far only been studied at the short-read level (Harr and Turner 2010).

In the present study, we explore the brain isoform diversity in house moue natural populations and closely related outgroup species. Using an optimized protocol to capture predominantly full-length capped transcripts, we identify widespread brain isoform diversity. We assess whether the individual isoform landscape can be used to trace the differentiation history of house mouse natural populations. We further analyze the origin of the isoform diversity and explore the possible source of genes and local alternative splicing events that contribute most to the overall isoform diversity. With the newly generated comprehensive transcriptomic data, we aim to define the set of conserved isoforms across mouse species, which are of particular relevance for functional studies that use the mouse as a biomedical model system in comparison to humans.

## Results

### An optimized experimental protocol to enrich full-length transcript

A key step for full-length RNA sequencing is the enrichment of intact transcripts without degradation, ideally with hallmarks of both, a 5’ CAP and a poly(A) tail. It has been reported that the TeloPrime full-length cDNA Amplification kit could selectively synthesize cDNA molecules from mRNAs carrying a 5’ CAP, in complement to the standard PacBio cDNA library preparation protocol that could only selectively synthesize cDNA molecules from transcripts with a poly(A) tail (Cartolano et al. 2016). In this study, we developed a new PacBio Iso-Seq library preparation protocol via combining 5’CAP capture in the TeloPrime full-length cDNA Amplification kit and poly(A) tail enrichment in the standard PacBio cDNA synthesis kit, which can synthesize cDNA molecules from transcripts with both a 5’ CAP and a poly(A) tail.

To test the performance of the TeloPrime-based protocol in an actual sample, we selected one mouse individual from the Taiwan population (Supplemental Data S1A) and implemented three distinct cDNA library preparation protocols for the RNA samples extracted from the brain (Figure 1A): i) standard PacBio cDNA library preparation protocol; ii) TeloPrime-based protocol with cDNA Amplification Kit v1; iii) TeloPrime-based protocol with cDNA Amplification Kit v2. The TeloPrime-based kit v2 includes updated reaction conditions compared to kit v1, while the method of cDNA generation remains unchanged. Each of the three cDNA libraries was sequenced on one SMRT cell of the same sequencing run, and the sequencing reads were analyzed and compared using the same computational approach (see Methods). Comparably high numbers of CCS reads were generated from each SMRT cell of the three kits (Supplemental Data S1A, Clontech: 472 k; Teloprime v1: 424 k; Teloprime v2: 437 k), ensuring the equivalent high-quality of these SMRT cells.

**Figure 1.**
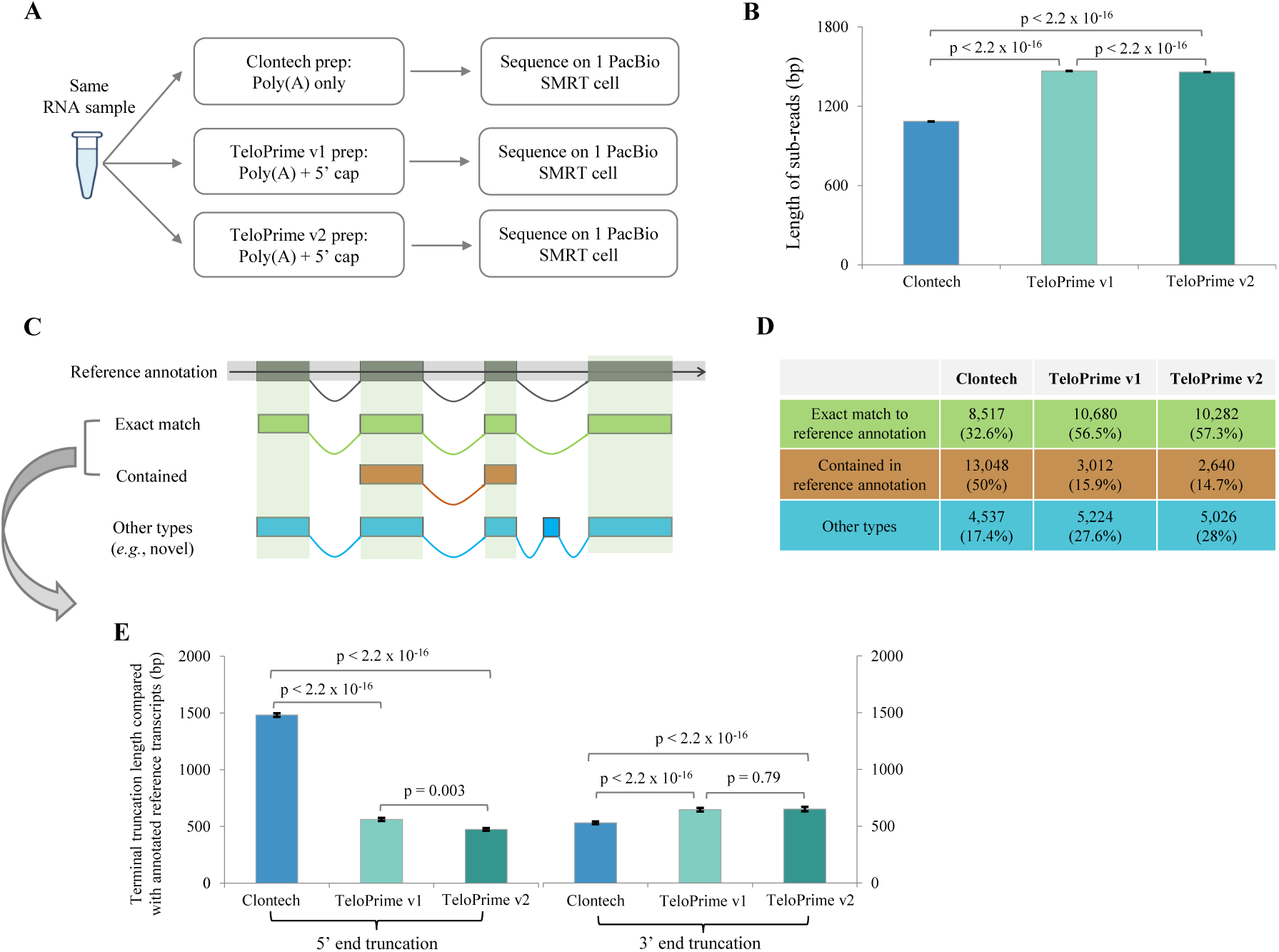
Performance evaluation of three types of full-length transcript enrichment protocols. (A) shows a simplified flow chart of this evaluation experiment setup. (B) indicates the length distribution of sub-reads generated from three PacBio Iso-Seq libraries. (C) illustrates the major categories of Iso-Seq transcripts based on their alignment coordinates to the reference genome, and (D) for the distribution of these categories. (E) shows the terminal truncation length of Iso-Seq transcripts from both 5’ and 3’ ends, compared to the reference annotation in Ensembl v103. Only Iso-Seq transcripts with exact matches or wholly contained in reference annotation were taken for analysis. The error bars indicate the standard error of means (SEM) of the length values. Statistical p-values were computed using two-sided Wilcoxon rank sum tests.

As shown in Figure 1B, we found that the sub-reads generated from TeloPrime-based cDNA libraries are, on average, longer (TeloPrime v1: 1,467bp; TeloPrime v2: 1,459bp) compared with those from the standard Clontech cDNA library (mean value: 1,085bp). Compared with that from the Clontech cDNA library (32.6%), a much higher fraction of isoforms from the TeloPrime v1 cDNA library (56.5%) and TeloPrime v2 cDNA library (57.3%) showed a complete and exact match of the exon chain as the annotated reference transcripts (Figure 1C and 1D). Moreover, around 50% of the isoforms from the Clontech cDNA library showed truncation of terminal exons. In comparison, this value is only 15.9% in the TeloPrime v1 cDNA library and 14.7% in the TeloPrime v2 cDNA library, suggesting that the TeloPrime-based cDNA enrichment protocols are more likely to enrich full-length transcripts. An in-depth investigation of the truncation lengths showed that the isoforms from the Clontech cDNA library are actually at least 2-fold more severely truncated from 5’end than those from TeloPrime-based cDNA libraries (similarly less extent from 3’end, Figure 1E), which confirmed the reliability of TeloPrime-based cDNA kits to selectively enrich mRNA transcripts carrying a 5’ CAP (Cartolano et al. 2016). Analysis of the 5’-exons of isoforms that were truncated in the Clontech kit showed that the 5’ terminal exons of the references transcripts tended to be longer (Supplemental Figure S1B) and more structured in an overall RNA structure analysis (Supplemental Figure S1C), both factors that may have contributed to the truncation tendency.

These findings suggest that the newly developed TeloPrime-based protocol provides a better approach to enriching full-length transcripts. Given the optimized reaction conditions, including the selection for capped RNA, we applied the TeloPrime-based protocol with cDNA Amplification Kit v2 to generate cDNA libraries for PacBio Iso-Seq for the following analyses.

### Alternative isoform landscape in wild mice brains

We analyzed the isoform landscape in whole brain transcriptomes for forty-eight unrelated outbred wild-type mice individuals raised under tightly controlled laboratory conditions (Supplemental Data S1B). They included forty house mouse (*Mus musculus*) individuals derived from five natural populations in the three major subspecies (*M. m. domesticus*, *M. m. musculus*, and *M. m. castaneus*), as well as eight individuals from two closely related outgroup species (*Mus spicilegus* and *Mus spretus*) (geographical origins and evolutionary relationships are depicted in Figure 2A). We performed both PacBio Iso-Seq and Illumina RNA-Seq for each brain sample to generate high-quality transcriptomes. As a result, an average of 60.1 (SD: 9.6) million raw sub-reads and 28.2 (SD: 3.3) million raw read pairs per sample were produced for PacBio Iso-Seq and Illumina RNA-Seq, respectively (Supplemental Table S1). This dataset represents the largest full-length isoform category at a comparative population level to date.

**Figure 2.**
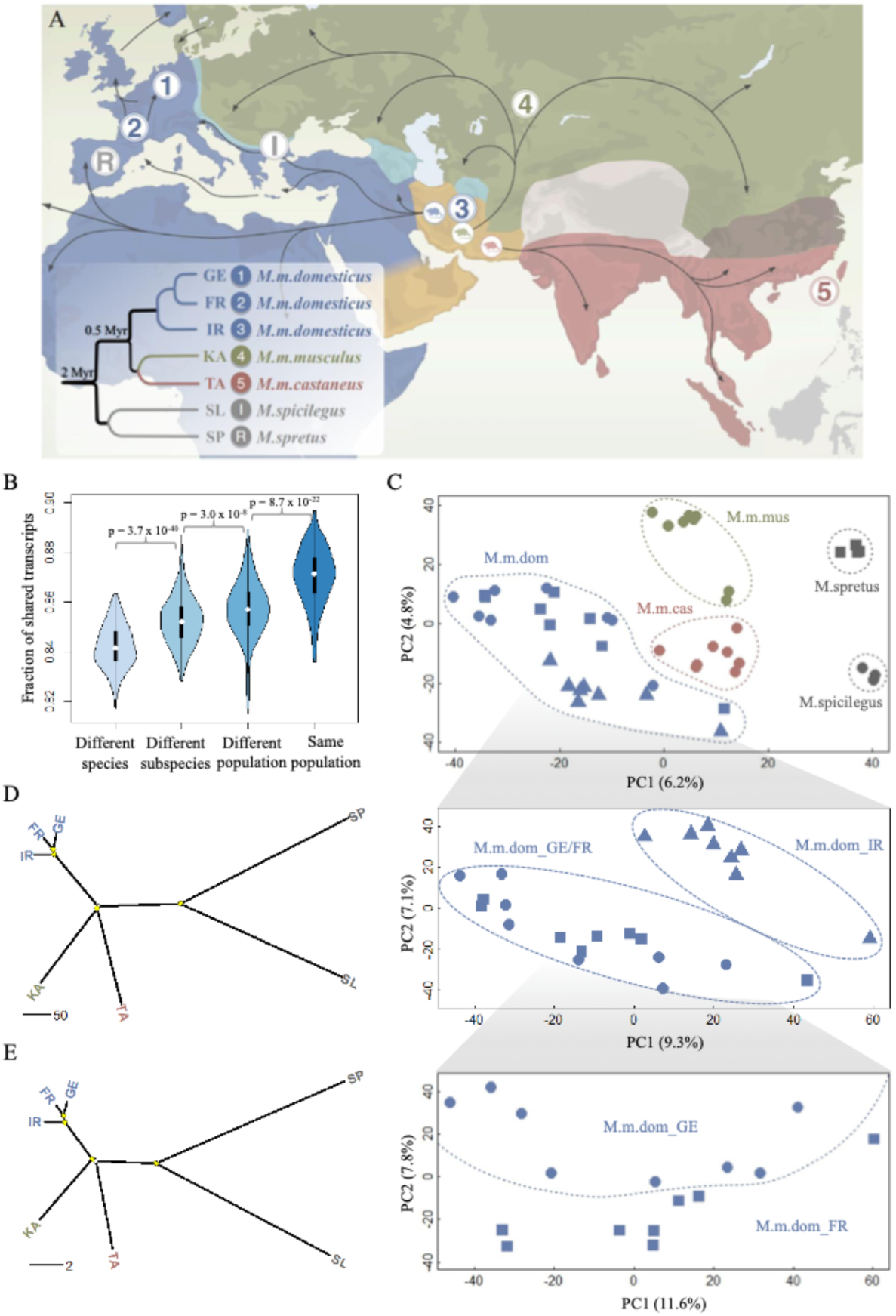
Geographic locations and phylogenetic relations of sampled mice individuals. (A) Geographic locations for the sampled mouse populations in this study. Territory areas for each house mouse subspecies: *M. m. domesticus* (blue), *M. m. musculus* (green), and *M. m. castaneus* (brown). Black arrows indicate possible migration routes, mainly during the spread of agriculture and trading. The inset figure shows the canonical phylogenetic relationships among the house mouse populations and outgroup species (branch lengths not scaled). Geographic locations: 1, Germany (GE); 2, France (FR); 3, Iran (IR); 4, Kazakhstan (KA); 5, Taiwan (TA). The sampling locations for the outgroup species are I, Slovakia (SL) for *M. spicilegus* and R, Spain (SP) for *M. spretus*. (B) shows the fraction of shared isoforms for pair-wise individual comparison. The fraction of shared isoforms between each pair of individuals is defined as the number of overlapping isoforms divided by the average number of detected isoforms of the two individuals to compare. Black boxes represent the interquartile range (IQR, distance between the first and third quartiles), with white dots in the middle to denote the median. The boundaries of the whiskers (also the ranges of violins) are based on the 1.5 IQR values for both sides. The statistical p-values of the fractions of shared isoforms between different comparison groups were computed using Wilcoxon rank sum tests. (C) shows the projection of the top two PCs of isoform variation in house mouse and outgroup individuals. Enlarged insets represent the results for the three populations from the subspecies of *M. m. domesticus*, as they cannot be well distinguished in the main figure. (D) and (E) show phylogenetic trees built based on isoform and SNP variants fixed within each population, respectively. Split nodes marked in yellow are the ones with bootstrap support value >70%.

An overview of the methodology to generate the high-quality transcriptome is given in Supplemental Figure S2. In brief, the raw PacBio Iso-Seq subread data were processed to produce circular consensus sequences (CCS) and further refined to generate full-length non-chimeric (FLNC) reads using the IsoSeq3 pipeline. To identify high-confidence isoforms, the FLNC reads were subject to *de novo* clustering using the same IsoSeq3 pipeline to generate an average of 49,147 (SD: 4,554) non-redundant isoforms per sample (Supplemental Data S1B), with the support from at least two FLNC reads (see below for the justification of this cutoff). We aligned the unique isoforms of each sample to the GRCm39/mm39 reference genome using minimap2 (Li 2018) and used the TAMA program (Kuo et al. 2020) with optimized parameters (Supplemental Text of Methods and Supplemental Figure S3) to collapse and merge the transcript models across all 48 samples into a single non-redundant transcriptome. We further refined a computational pipeline to filter out low-quality transcripts and those of potential artifacts (see Methods), resulting in a total of 117,728 high-confidence distinct isoforms derived from 15,012 distinct loci (Supplemental Data S2). For reference, we also include the list of isoforms covered only by singleton reads (Supplemental Data S3), but we do not include these in the further analyses.

To evaluate the reliability of these isoforms, we assessed their quality for three distinct criteria separately: i) transcription start site (TSS), ii) transcript end site (TES), and iii) splice junctions (SJ). Reliable TSSs are defined as those supported by Ensembl GRCm39/mm39 reference genome annotation (Howe et al. 2021), collection of experimentally validated TSS peaks (Abugessaisa et al. 2019), and a significantly greater coverage ratio between inside isoform and outside isoform than the null hypothesis (Tardaguila et al. 2018) (Supplemental Figure S4). Similarly, reliable TESs are defined as those supported by the reference genome annotation (Howe et al. 2021) and collection of experimentally validated poly(A) sites and motifs (Herrmann et al. 2020). The reliable SJs are defined as those present in reference genome annotation (Howe et al. 2021) and those belonging to the canonical splicing signals (GT-AG, GC-AG, and AT-AC) (Parada et al. 2014). A more detailed discussion on evaluating the reliability of the detected isoforms can be found in the Supplemental Text of Methods, including Supplemental Figures S5 and S6. Based on the above-defined criteria, we found that 98.7% of the TSSs, 94.4% of the TESs, and 99.1% of the SJs from all the detected isoforms are supported by respective previous evidence (Supplemental Data S2), and a comparable large extent (>96% for all three features) of isoforms could be validated for all the seven tested populations (Supplemental Figure S7).

### The role of splicing noise

Since biochemical systems are never perfect, one should expect a certain number of errors in generating isoforms (Pickrell et al. 2010; Wan and Larson 2018). This could be considered as noise rather than being regulated through genetic polymorphisms or epigenetic effects.

Noise should be non-deterministic in the sense that the absolute number of isoforms generated by noise should increase with increased read sampling. This can be tested by a rarefaction analysis, where one takes increasingly larger random subsets of the reads in a given experiment and asks whether the number of isoforms keeps increasing. This is indeed the case. When using all reads for a given individual, one gets a linear increase in the number of isoforms with increased sampling without any signs of saturation (Figure 3A). However, when one does the same analysis for the high-confidence isoforms represented by at least 2 FLNC reads in a given individual, one gets an almost perfect saturation at the sequencing depth applied in our experiments (Figure 3B). The same patterns can be seen for individuals from each population sequenced (Supplemental Figures S8 and S9). Because of this difference in saturation behavior, we conclude that singletons are mostly the product of splicing errors, consistent with conclusions in previous studies (Pickrell et al. 2010; Saudemont et al. 2017).

**Figure 3.**
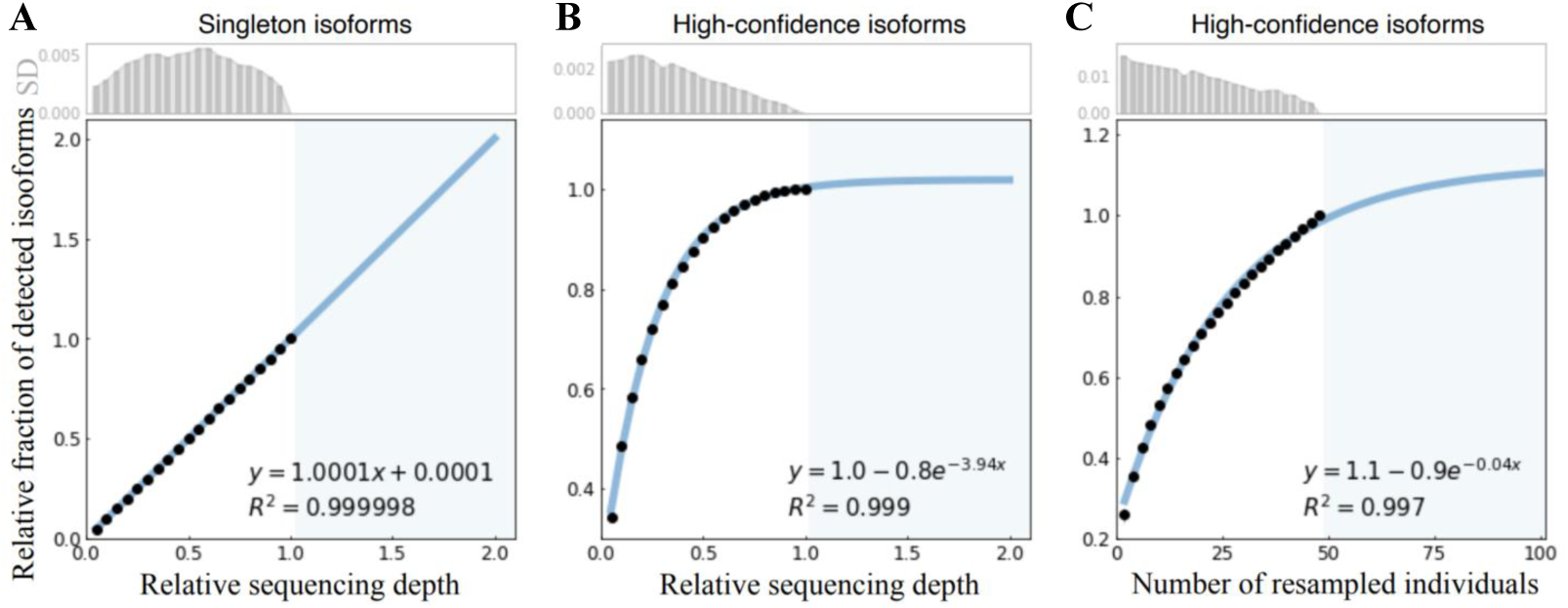
Saturation analysis on the sequencing depth and number of sampled individuals. (A) and (B) show the relative fractions of detected isoforms supported with singleton FLNC read versus two or more FLNC reads (high-confidence isoforms) with increasing random resampling of the Iso-Seq sequencing depth, respectively. The resampling sequencing depths were selected from 0.05 to 1, with a step size of 0.05. The blue area shows the prediction after doubling the actual Iso-Seq sequencing depth. The illustration is based on one randomly selected individual (GE3) in the GE population; the analogous results for individuals from other populations are provided in Supplementary Figures S8 and S9. (F) Relative fractions of detected isoforms with increasing random resampling sample sizes of individuals in the main experiment. The resampling sequencing sizes were selected from 2 to 48, with a step size of 2. The blue area shows the prediction after doubling the current sampling of mice individuals. The bar plots show the standard deviation (SD) of each resampling analysis.

Interestingly, random resampling analysis of subsamples across individuals shows that the number of detectable high-confidence isoforms remains unsaturated with the number of sampled individuals in our dataset (Figure 3C). That shows that more new isoforms are expected to show up when more individuals are analyzed, suggesting that these additional isoforms are more likely generated by genetic polymorphisms between the individuals than by noise. However, some saturation should eventually be expected even at this level since there is a limit to the maximum number of isoforms that can be reliably maintained in natural populations (Benitiere et al. 2024).

In a further analysis of the noise effect, we make use of a recent discovery on the role of transposable elements (TE) influencing splicing. Ilik et al. (Ilik et al. 2024) found that cells have developed mechanisms to maintain splicing integrity with respect to excluding cryptic splice sites generated by TEs. If this process is leaky, isoforms generated by noise effects should also include more TEs in their exons. We find that this is indeed the case. Comparison of the transposon element (TE) insertions in singleton isoforms versus high-confidence isoforms shows that the latter harbor a significantly lower fraction of TE fragment insertions (Fisher’s exact test, p-value < 2.2 x 10^-16^, Supplementary Figure S10).

Based on these analyses, we concluded that by excluding the singleton reads from the further analyses, we are indeed excluding most of the effects of noise. Evidently, it is still possible that reads that occur more than once are generated through noise effects due to the chromatin context in which they are transcribed or special structures of their RNAs. However, we consider the set of high-confidence isoforms to be at least highly enriched in variants that are not simply generated by errors in the splicing machinery.

### Alternative isoforms as population differentiation markers

Given the high and comparable numbers of isoforms in wild mice brain transcriptomes across populations (Table 1), we asked whether the isoform diversity landscape can distinguish individuals, *i.e.*, whether they could contribute to individual differences. If this is the case, they could also be used to trace the divergence trajectories of the populations.

**Table 1.** Statistics on the detected isoforms in the house mouse populations and outgroups. The digit succeeding the character N indicates the number of sequenced mouse individuals in each population. The graphic depiction of isoform categories is shown in Figure 4A. Isoforms categorized as FSM and ISM are defined to be known isoforms, gene loci with any isoform not categorized as either Antisense or Intergenic to be annotated genes. SD values refer to variances between the individuals in a given population.

|  | *M. m. domesticus* |  |  | *M. m. musculus* | *M. m. castaneus* | *M. spicilegus* | *M. spretus* |
| --- | --- | --- | --- | --- | --- | --- | --- |
|  | GE (N: 8) | FR (N: 8) | IR (N: 8) | KA (N: 8) | TA (N: 8) | SL (N: 4) | SP (N: 4) |
| **Unique isoforms** | 23,027 (SD:2,001) | 21,609 (SD:2,315) | 21,763 (SD:2,006) | 22,380 (SD:1,553) | 21,279 (SD:1,532) | 20,644 (SD:858) | 21,640 (SD:640) |
| Protein-coding | 21,394 (SD:2,062) | 20,024 (SD:2,431) | 20,155 (SD:1,941) | 20,798 (SD:1,486) | 19,642 (SD:1,435) | 18,786 (SD:929) | 20,063 (SD:597) |
| Nonsense-mediated decay | 1,152 (SD:226) | 1,071 (SD: 220) | 1,098 (SD:209) | 1,125 (SD:146) | 1,142 (SD:216) | 1,038 (SD:146) | 1,131 (SD:46) |
| Non-protein-coding | 1,634 (SD:239) | 1,585 (SD:284) | 1,608 (SD:324) | 1,582 (SD:235) | 1,637 (SD:261) | 1,858 (SD:73) | 1,577 (SD:63) |
| **Known isoforms** | 17,266 (SD:1,216) | 16,306 (SD: 1,419) | 16,343 (SD:1,257) | 16,487 (SD:861) | 15,765 (SD:821) | 15,202 (SD:512) | 15,608 (SD:416) |
| Full Splice Match (FSM) | 15,007 (SD:1,217) | 14,245 (SD: 1,441) | 13,654 (SD:1,053) | 14,466 (SD:810) | 13,561 (SD:602) | 12,812 (SD:523) | 13,621 (SD:297) |
| Incomplete Splice Match (ISM) | 2,259 (SD:297) | 2,060 (SD: 654) | 2,689 (SD:531) | 2,021 (SD:593) | 2,204 (SD:313) | 2,389 (SD:325) | 1,988 (SD:154) |
| **Novel isoforms** | 5,762 (SD:796) | 5,304 (SD: 913) | 5,421 (SD:778) | 5,893 (SD:708) | 5,514 (SD:768) | 5,442 (SD:415) | 6,032 (SD:237) |
| Not In Category (NIC) | 3,765 (SD:599) | 3,490(SD:651) | 3,478 (SD:514) | 3,611 (SD:459) | 3,469 (SD:528) | 3,204 (SD:318) | 3,582 (SD:165) |
| Novel Not in Category (NNC) | 1,811 (SD:202) | 1,647 (SD: 271) | 1,768 (SD:256) | 2,066 (SD:254) | 1,839 (SD:234) | 2,015 (SD:120) | 2,248 (SD:59) |
| Fusion (F) | 74 (SD:11) | 67 (SD:15) | 78 (SD:11) | 90 (SD:18) | 93(SD:17) | 65 (SD:7) | 84 (SD:8) |
| Genic (G) | 22 (SD:5) | 19 (SD:4) | 18 (SD:5) | 22 (SD:4) | 25 (SD:4) | 24 (SD:1) | 24 (SD:3) |
| Antisense (A) | 35 (SD:8) | 30 (SD:10) | 33 (SD:12) | 32 (SD:6) | 30 (SD:10) | 49 (SD:12) | 31 (SD:3) |
| Intergenic (I) | 54 (SD:7) | 51 (SD:13) | 44 (SD:14) | 72 (SD:10) | 59 (SD:15) | 84 (SD:6) | 62 (SD:10) |

A pairwise individual comparison of the sharing of isoforms reveals an increasingly larger fraction of shared isoforms for individuals from different subspecies, different subspecies but same species, different populations but same subspecies, and same populations (Figure 2B). The fraction degrees of shared isoforms between different comparison groups are all statistically significant (p-value < 0.05). The overall principal component analysis (PCA) shows clear clusters of individuals according to their origin. Still, considerable diversity within each cluster implies that isoform diversity indeed confers a form of individuality to each animal (Figure 2C). We further used the presence/absence matrix of fixed isoforms within each population to build an overall phylogeny (Figure 2D), which has the same general topology as a phylogeny based on the SNP variants called from Illumina RNA-Seq datasets from the same set of mice individuals (Figure 2E).

The above patterns were unlikely caused by the imbalance of sequencing depth for different mice individuals, as the numbers of detected isoforms have reached saturation at the individual level (Figure 3B). These findings remain valid when analyzing the isoforms observed in at least two individuals (Supplemental Figure S11), further eliminating the possible dominating effects of low-abundance isoforms.

Given these observations, we performed an isoform frequency-based approach to detect the isoforms as population-differentiating markers (see Methods). Each isoform was assigned a population differentiation index (PDI) to indicate its power to differentiate populations. After applying a false discovery rate (FDR) threshold of 0.05, we detected in total 113,008 isoforms (96% of all isoforms) that could differentiate populations from house mouse and outgroup species (with significantly larger PDI values compared with null distribution from 1,000 times random shuffling; see Supplemental Data S2). We conclude that isoforms can serve as effective markers to differentiate house mouse natural populations.

### A high number of novel isoforms are absent in the reference annotation

To further characterize the features of the 117,728 high-confidence distinct isoforms, we compared them with those annotated in the GRCm39/mm39 reference genome from Ensembl v103 (Howe et al. 2021), built largely based on a single C57BL/6 lab mouse inbred strain, which has predominantly a genomic *M. m. domesticus* population background. Based on their alignment status to the Ensembl mouse transcriptome v103 (Howe et al. 2021), these isoforms were classified into eight distinct structural categories using SQANTI3 (Tardaguila et al. 2018) (Figure 4A): i) full splice match (FSM); ii) incomplete splice match (ISM); iii) novel in category (NIC); iv) novel not in category (NNC); v) Fusion (F); vi) genic (G); vii) antisense (A); viii) intergenic(I). The statistics on the number of isoforms from each structural category at the population level can be found in Table 1.

**Figure 4.**
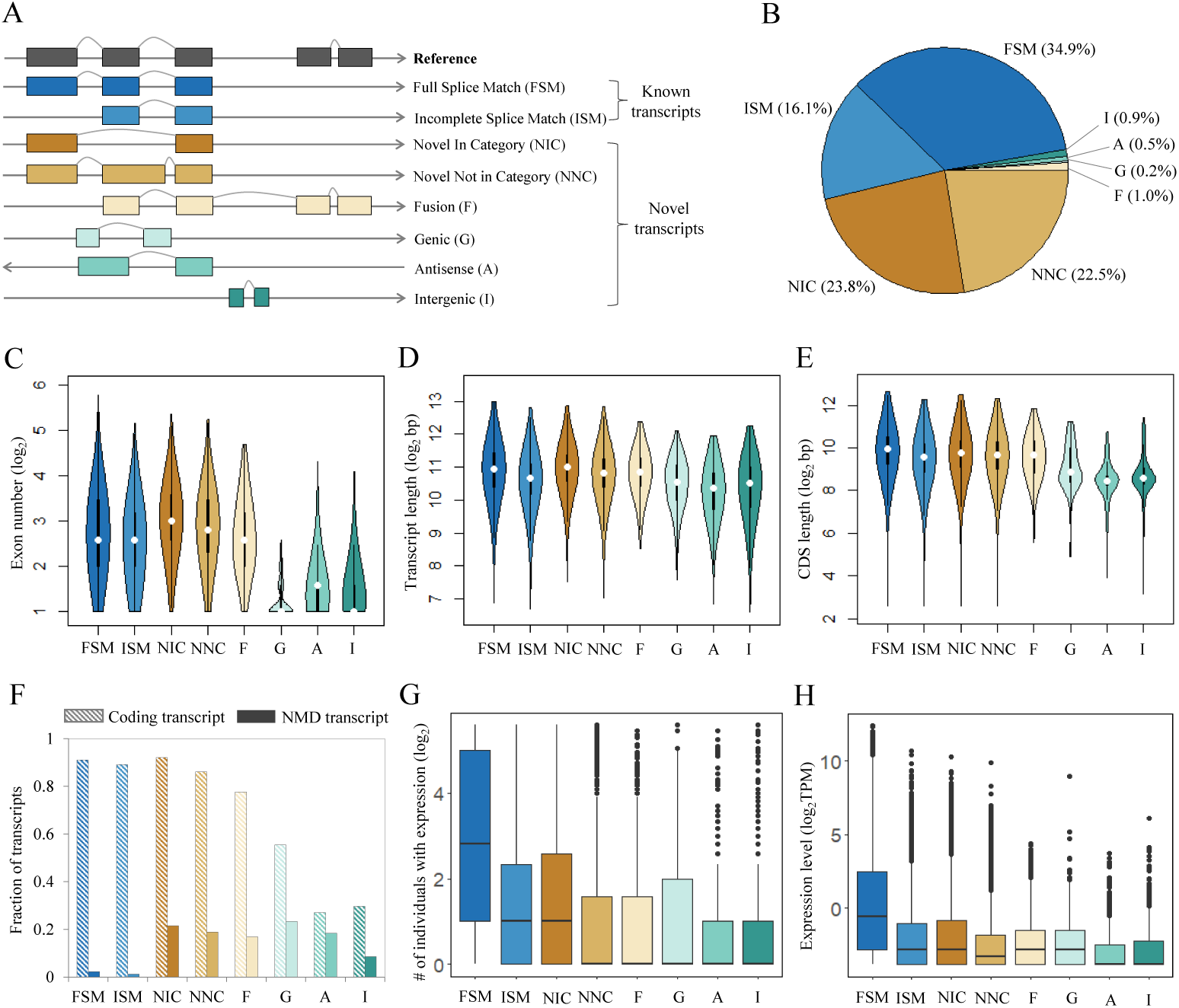
Characterization of all detected high-confidence isoforms. (A) Types and illustrations of identified isoforms. (B) Fraction distribution of isoform structural categories (see isoform annotations in Supplemental Data S2). (C)-(H) show the distributions of isoform types for distinct features. Isoforms with expression were defined as those with non-zero TPM values, and the expression levels were computed based on the number of supported FLNC reads using SQANTI3 (Tardaguila et al. 2018). Boxes represent the interquartile range (IQR, distance between the first and third quartiles), with white dots (or black lines) in the middle to denote the median. The boundaries of the whiskers (also the ranges of violins for panels C-E) are based on the 1.5 IQR values for both sides; black dots in G and H represent outliers.

In total, we found only 51% of the 117,728 distinct isoforms matching perfectly to a whole (FSM, 34.9%) or subsection (ISM, 16.1%) of a reference annotated isoform, designated as known isoforms following the convention in (Leung et al. 2021; Naftaly et al. 2021). The remaining 49% (57,730) of the identified isoforms were missing in the Ensembl transcriptome (Figure 4B), including the ones deriving from the rearrangement and modification of annotated exons (novel combination of known splice sites or NIC: 23.8%; novel splice sites or NNC: 22.5%; and merging of adjacent transcripts or Fusion: 1.0%), from intronic regions (Genic: 0.2%), and from entirely novel gene loci (Antisense: 0.5%; Intergenic: 0.9%). Of note, the novel isoforms exhibit comparable high quality and reliability as the known isoforms, based on the confirmation of the transcription start site, splice junction, and transcript end site (Supplemental Figure S5C-E).

One may argue that the isoforms specific to outgroup species could contribute substantially to the novel isoform category, as the GRCm39/mm39 reference genome was built primarily based on a single C57BL/6 lab inbred strain (*i.e.*, *Mus musculus*). The individuals from the outgroup species indeed have an average of 27.1% of isoforms that are not registered in the mm39 reference annotation, slightly larger than the value of 25.2% in mice individuals belonging to *Mus musculus* (p-value = 0.003, Wilcoxon rank sum test, Supplemental Data S4). However, we still found that around 43% of the identified unique isoforms were missing in the reference annotation after excluding those isoforms detected only in individuals of outgroup species (Supplemental Figure S12B). Of note, the fraction of FLNC reads to support novel isoforms is generally lower than the fraction of novel isoforms for each mouse individual (Average value: 6.0% vs 25.5%, Supplemental Data S4).

Given that the current reference annotation is based on an isogenic strain, it represents nominally only the isoforms of a single individual but not the isoforms typical for a population or across populations. At the same time, it includes isoforms that are restricted to its population of origin. We have therefore compiled a set of 39,537 isoforms that are conserved in the house mouse and the outgroup species (marked in yellow in Supplemental Data S2). A substantial fraction (30.6%) of these isoforms is not included in the reference annotation (Supplemental Figure S12C). We propose that this conserved set of isoforms should be the best reference when comparing between mice and other species, *e.g.*, humans.

In comparison to known isoforms (FSM, ISM), novel isoforms deriving from annotated exonic regions (NIC, NNC, and F – which constitute the bulk of the new isoforms) show comparable exon numbers (Figure 4C), transcript length (Figure 4D) and CDS length (Figure 4E). The novel isoforms from intronic (G) and unannotated gene loci (A, I) show generally lower values for all these features (Supplemental Table S2). Notably, all the novel isoforms with coding potential have a significantly higher probability of becoming degraded via the nonsense-mediated decay (NMD) process (Chang et al. 2007) due to premature translation-termination codons (PTCs – detected by SQANTI3) than known isoforms (Figure 4F, mean 0.18 vs. 0.02, Fisher’s exact test, p-value < 2.2 x 10^-16^). Most novel isoforms were found to be more restrictively expressed in a smaller number of individuals, except for NIC, which are comparable to ISM in this respect (Figure 4G). The general expression levels of the novel isoforms, except A and I, are at a similar level as the known ISM isoforms (Figure 4H). Regarding absolute isoform abundance, the newly discovered isoforms contribute about 6% to the total number of expressed isoforms in the overall dataset, in contrast to 49% of identified unique isoforms as novel ones (Figure 4B).

### Housekeeping genes are the major contributor to isoform diversity

To analyze the characteristics of the gene loci that contribute significantly to the overall isoform diversity, we classified all the detected gene loci into three roughly similar-sized classes based on their splicing isoform numbers (Figure 5A): i) Few-isoform Genes (FG, with no more than two isoforms, n = 5,372), ii) Many-isoform Genes (MG, with isoform number between three and eight, n = 5,012), and iii) Plenty-isoform Genes (PG, with no less than nine isoforms, n = 4,628). Among all the 117,728 high-confidence isoforms detected in the wild mice brain transcriptome, 4.3% are derived from the FG group, 18.2% from the MG group, and the largest fraction (77.4%) from the PG group.

**Figure 5.**
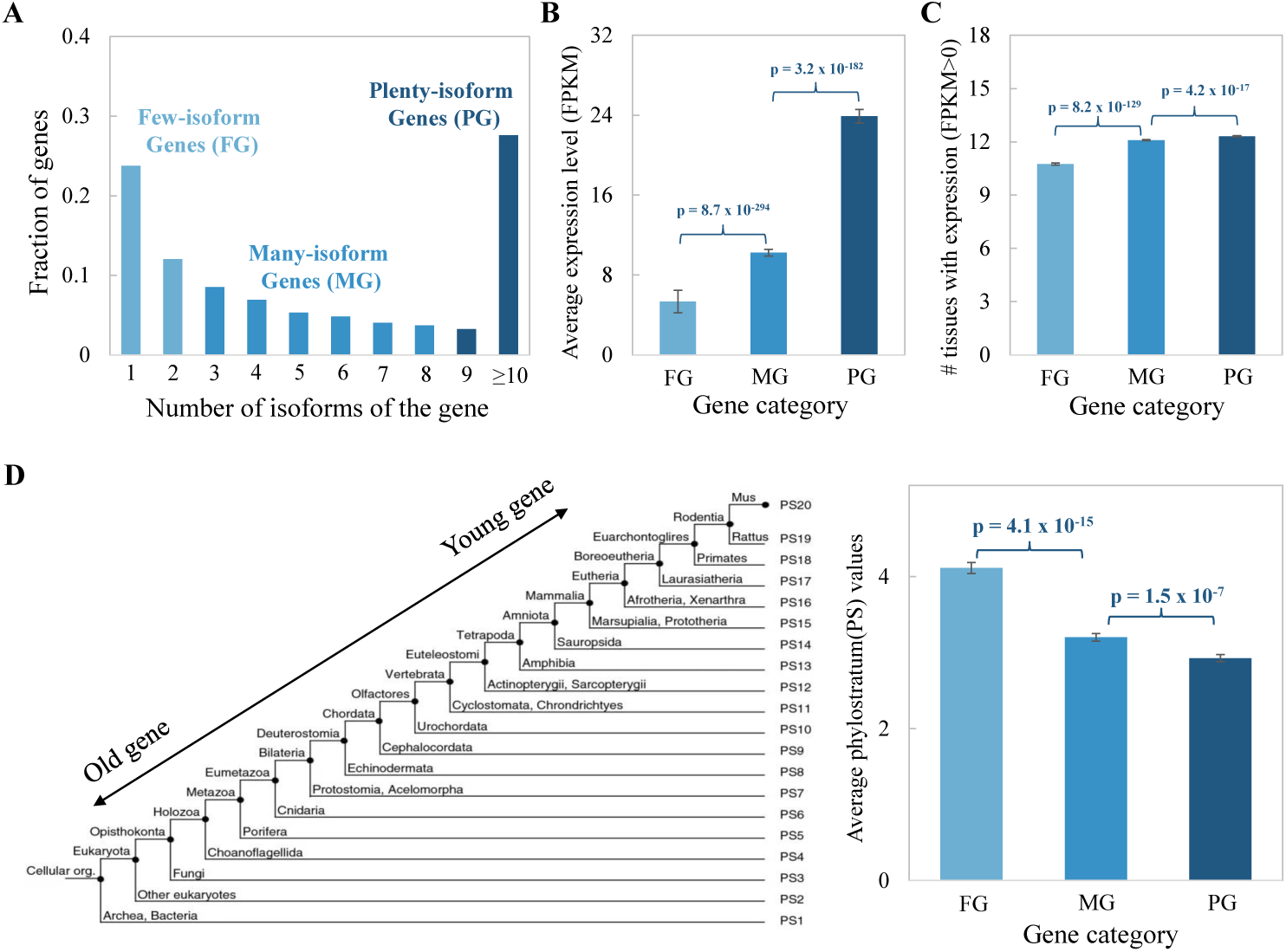
Characteristics of gene loci with different numbers of isoforms. (A) Gene categories were based on the number of isoforms of each gene. Three gene categories were classified with roughly the same number of genes in each group. (B) Comparison of the average gene expression level among the three gene categories. The average gene expression level was calculated based on the Illumina RNA-Seq generated in this study. (C) Comparison of the number of tissues with expression among the three gene categories. The transcriptomic sequencing data from 13 tissues of the *Mus musculus* C57BL/6 inbred line was retrieved from (Lin et al. 2014). The detectable gene expression in each tissue was defined as FPKM > 0. (D) Comparison of the average phylostratum (PS) values among the three gene categories. The PS assignment data for mouse protein-coding genes were retrieved from (Neme and Tautz 2013). The smaller PS values indicate the older gene groups. Error bars show standard errors of the mean (SEM) values of average gene expression levels, number of tissues with expression, and average PS values, respectively. Statistical p-values were computed using two-sided Wilcoxon rank sum tests.

Based on the gene expression analysis of matched Illumina RNA-Seq dataset, we found that genes with more splicing isoforms have a significantly higher expression level in the brain (Figure 5B, MG vs. FG: p-value = 8.7 x 10^-294^, PG vs. MG: p-value = 3.2 x 10^-182^, two-sided Wilcoxon rank sum tests). Additionally, through analyzing the transcriptomic sequencing dataset from thirteen tissues of the *M. m. domesticus* derived C57BL/6 inbred line (Lin et al. 2014) (Supplemental Data S1D), we observed a clear trend that genes with more splicing isoforms tend to be expressed significantly more broadly, *i.e.*, with expression in more tissues (Figure 5C, MG vs. FG: p-value = 8.2 x 10^-129^, PG vs. MG: p-value = 4.2 x 10^-17^, two-sided Wilcoxon rank sum tests). Furthermore, gene ontology (GO) enrichment analysis on the PG group showed that they are involved in a variety of biological processes for the maintenance of cellular function (Supplemental Data S5), a common feature of housekeeping genes (Eisenberg and Levanon 2003). These findings suggest that housekeeping genes contribute most to the overall isoform diversity.

Previous studies have shown that evolutionarily old genes usually have a broader tissue expression and higher expression levels (Zhang et al. 2015). Thus, it can be hypothesized that genes with more splicing isoforms tend to be evolutionarily older. We set out to test this hypothesis via a comparative analysis of the gene age distribution pattern among the gene groups with different numbers of splicing isoforms, for which the mouse gene age assignment data was retrieved from (Neme and Tautz 2013). Our analysis indeed showed that there is a significant enrichment in evolutionarily older gene groups for those genes with more splicing isoforms (Figure 5D, MG vs. FG: p-value = 4.1 x 10^-15^, PG vs. MG: p-value = 1.5 x 10^-7^, two-sided Wilcoxon rank sum tests).

### Isoform expression in highly versus lowly expressed genes

It has previously been noted that the fraction of isoforms expressed as alternative isoforms from highly expressed genes is lower than that for lowly expressed genes (Sharon et al. 2013; Benitiere et al. 2024). We have analyzed this in our data by generating normalized FLNC read counts (TPM) as a proxy for the expression level of the respective isoforms from each gene. We then compared the top expressed (T) isoform for a given gene with the read counts of the remaining isoforms (R) from the same gene. We restrict this analysis to *M. musculus* populations, *i.e.*, excluding the outgroups, since many top expressed isoforms change in these outgroups. The average ratio R/T across all genes with at least two isoforms in at least one individual of the *M. musculus* populations (N = 9,111) is 1.03 (SD: 1.04). The same calculation for the top 10% expressed genes is 0.43 (SD: 0.63) and for the lowest 10% expressed genes is 1.11 (SD: 0.62) (Supplemental Data S6), *i.e.*, the overall fraction of alternative isoform molecules is about 3-fold lower for highly expressed genes compared to the lowest expressed genes (p-value = 2.02 x 10^-147^, Wilcoxon rank sum test). Hence, while highly expressed genes have, on average, more alternative isoforms than lowly expressed genes (see above), their relative expression level is much lower than for the alternative isoforms of lowly expressed genes. This could suggest constraints on highly expressed genes for generating too many new isoforms (Saudemont et al. 2017; Benitiere et al. 2024).

### Local alternative splicing events contribute to isoform diversity

The full-length isoforms appear as a combination of different types of local alternative splicing (AS) events. Thus, it is useful to disentangle the relative contribution of each type of AS event to the overall isoform diversity. For this purpose, we used the SUPPA2 program (Trincado et al. 2018) to identify different types of splicing events in the complete set of high-confidence isoforms, including skipped exon (SE), retained intron (RI), alternative 5’ splice site (A5), alternative 3’ splice site (A3), mutually exclusive exon (MX), alternative first exon (AFE), and alternative last exon (ALE) (Figure 6A).

**Figure 6.**
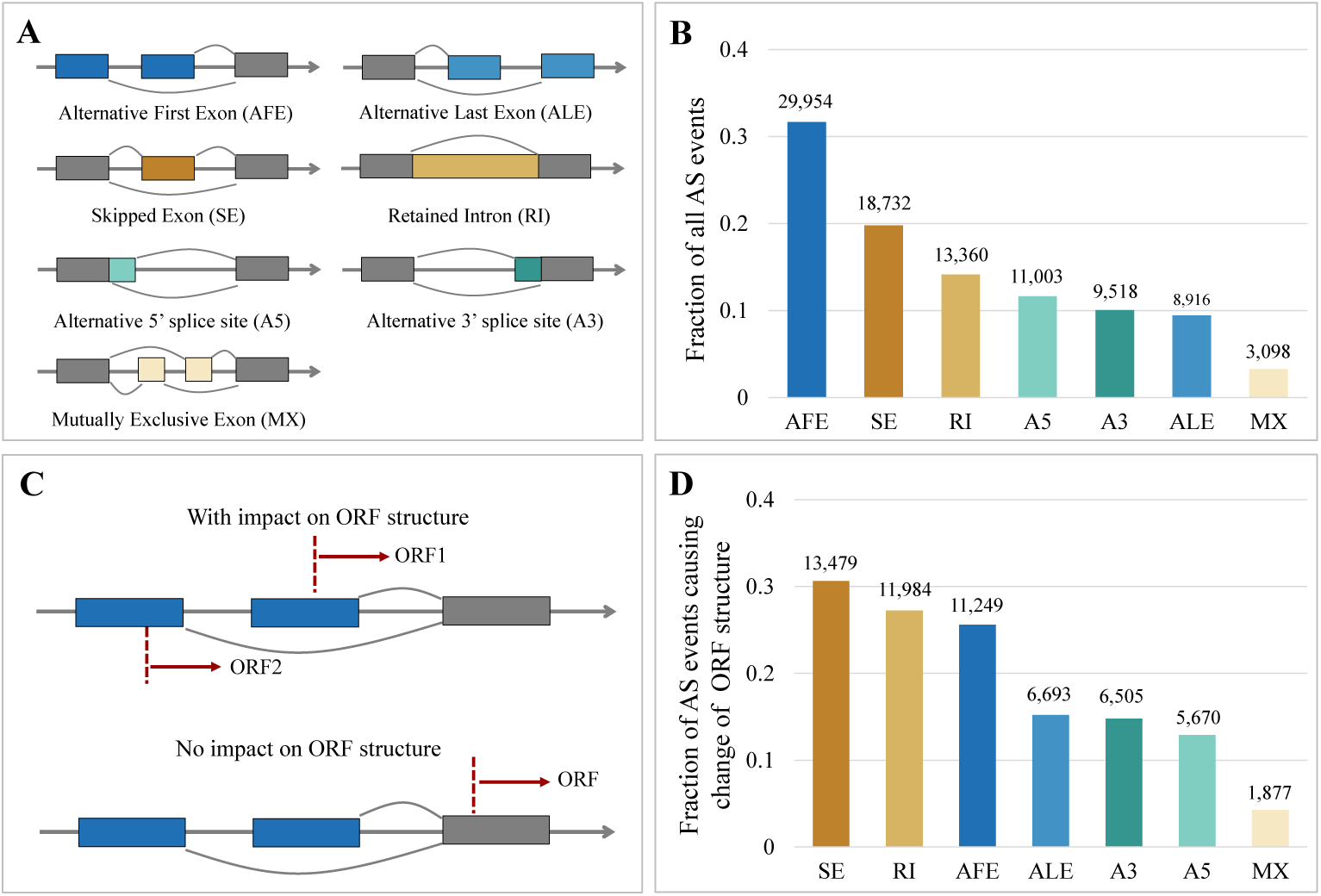
Distribution of different types of local AS events. (A) Types and illustrations of AS local events. (B) The distribution of all types of local AS events. (C) An example of AFE events that change ORF structures and similar situations for other types of local AS events. The dashed red lines indicate the in-frame start codon positions. (D) The distribution of all types of local AS events that impact respective ORF structures. The value above each bar in (B) and (D) indicates the number of respective types of local AS events.

Among all the 94,581 splicing events detected in all the isoforms (Figure 6B), AFE events contribute most to the overall isoform diversity (31.7%), followed by SE events (19.8%), RI events (14.1%), A5 (11.6%), A3 (10.1%), ALE (9.4%), and with MX as a minor contributor (3.3%). This finding is in line with the previous report on the AFE as the most prevalent splicing event for the overall transcriptome in the inbred laboratory mouse cerebral cortex (Leung et al. 2021), suggesting a dominant role of using AFE to generate alternative isoforms in mice brain transcriptomes.

AFEs would not necessarily impact the coding sequences (Landry et al. 2003). To address this issue, we performed an additional analysis by collapsing the isoforms with the same coding sequence (*i.e.*, only isoforms with predicted ORFs were considered) into a single unique ORF. We indeed observed a much lower number (less than two-thirds) of unique ORFs than isoforms in each mouse individual (Supplemental Figure S13). Based on the landing position of the start codon in relation to local AS events (Figure 6C), we further enumerated the number of local AS events causing the change of the respective coding sequences. We found that only 46.5% (43,978) of all the local splicing events impact the ORF structures (Figure 6D). Compared to AFE, SE, and RI events are more prevalent among all the local AS types to contribute to ORF diversity. The majority of AFE events (55%) would cause no change in the coding sequences while acting as a major source to generate isoform diversity at the RNA level (Huang et al. 2021). These data illustrate the distinct roles of local AS events in contributing to isoform and ORF diversity in wild mice brain transcriptomes.

## Discussion

We used PacBio Iso-Seq and Illumina RNA-Seq to characterize the full-length isoform diversity of the brain transcriptomes in house mouse natural populations, resulting in the first and most comprehensive full-length isoform category representation at a comparative population level to date. Via implementation of a separate 5’ CAP selection step (Cartolano et al. 2016; Kuo et al. 2017), our optimized approach improved the performance to enrich genuine full-length transcripts.

Our overall results confirm the conclusions from previous studies that the diversity of alternatively spliced isoforms surpasses the current annotation level, even of exceptionally well-curated genomes, such as the one from the mouse (Leung et al. 2021). Our study finds double as many isoforms as are currently annotated, suggesting the widespread diversity of alternative isoforms in natural populations. Intriguingly, these isoforms could be efficient markers to infer population relationships and resolve the phylogenetic tree of house mouse natural populations and outgroups. This suggests that the isoform diversity reflects directly the genetic polymorphisms segregating in these populations. Note that the mice used for this study were reared under tightly controlled laboratory conditions to eliminate possible environmental influences on isoform diversity.

Interestingly, evolutionarily old housekeeping genes with overall high expression in many tissues show the largest number of new splice variants on average, likely due to the longer gene length and a larger number of exons for the evolutionarily old housekeeping genes (Zhang et al. 2015). Indeed, we found plenty-isoform genes are significantly longer (indicated by the longest transcript length) and have significantly more exons (Supplemental Figure S14), potentially allowing more space for alternative splicing machinery to act on.

The focus on full-length transcripts revealed that new 5’-exons are the relatively most abundant source of new isoforms, followed by the “skipped exon” and “retained intron” variants, which were previously thought to be the most prevalent ones (Grau-Bove et al. 2018). Many “alternative first exon” isoforms might originate from new promotors that can easily develop upstream of existing genes. This is in line with the realization that enhancers as regulatory elements can also assume promotor functions (Andersson and Sandelin 2020). Only about half of the new splice variants affect the annotated ORF of the respective genes, implying that the ORF diversity generated through alternative splicing is lower than the one at the isoform level. However, new short ORFs or alternative ORFs that may arise in the alternatively spliced isoforms are yet to be studied (Leong et al., 2022).

Our dataset allows to define isoforms that should be of particular interest for functional comparisons across larger evolutionary distances, including humans. The currently annotated mouse genome also includes the isoforms that are population-specific for the subspecies from which the C57Bl/6 was derived. Most of these will have specifically arisen in this lineage and are, therefore, of less interest for broader functional comparisons. With our comparative data, however, we can define the set of isoforms that are typically found across multiple mouse species, including new ones that were so far missing from the annotation. We have compiled this more informative and more inclusive set of isoforms (Supplemental Data S7), which will be of particular use for broader comparative studies between mice and other species.

## Methods

### Ethics statement

All the mice were kept according to FELASA (Federation of European Laboratory Animal Science Association) guidelines, with the permit from the Veterinäramt Kreis Plön: 1401-144/PLÖ-004697. The samples used for this study were derived as organ retrieval under permit number 1158 (following §4 German TSchVersV) from the respective animal welfare officers at the University of Kiel (Prof. Schultheiss and Mrs. Vieten).

### Sample collection and RNA extraction

49 male mice individuals were sampled in this study, including one being used in the Iso-Seq library preparation protocol optimization experiment (Supplemental Data S1A and S1B). Eight individuals were chosen for each of the following populations covering all three major subspecies of the house mouse (*Mus musculus*): Germany, France, and Iran populations from *Mus musculus domesticus*, Kazakhstan population from *Mus musculus musculus*, and Taiwan population from *Mus musculus castaneus*. We also included four individuals from each of the two outgroup species, *Mus spretus* and *Mus spicilegus*. All mice were derived from previously wild-caught founder mice and were maintained in an outbred scheme under standard rearing techniques, which ensure a homogeneous environment (Harr et al. 2016). Each mouse individual was obtained from one distinct, unrelated breeding family. Thus, the relatedness between individuals, especially from the same population, should be minimized (see Supplemental Data S1E for pairwise relatedness estimates).

Mice were sacrificed at approximately ten weeks of age by CO_2_ asphyxiation followed immediately by cervical dislocation. The whole brain was dissected and immediately frozen in liquid nitrogen within 5 minutes post-mortem. Total RNAs were extracted and purified using RNeasy lipid tissue kits (Qiagen, The Netherlands). RNA was quantified using Qubit Fluorometers (Invitrogen, Thermo Scientific, USA), and RNA quality was assessed with 2100 Bioanalyzer (RNA Nanochip, Agilent Technologies, USA). All samples with RIN values above 8.5 were then used for both PacBio and Illumina transcriptome sequencing at the Max Planck-Genome-Centre Cologne.

### Selection of cDNA library enrichment protocol for PacBio Iso-Seq

It has been reported that the TeloPrime full-length cDNA Amplification kit, which selectively synthesizes cDNA molecules from mRNAs carrying a 5’ CAP in comparison to the standard PacBio cDNA library preparation protocol, provides a better solution to enrich for actual full-length transcripts (Cartolano et al. 2016). In this study, we selected one male mouse from the Taiwan population (Supplemental Data S1A) to test the performance of TeloPrime-based protocols in the real world.

We applied three distinct cDNA library enrichment protocols for the RNA sample extracted from the same mice individual: i) standard PacBio Clontech SMARTer PCR cDNA Synthesis kit (Clontech Laboratories, Inc.); ii) TeloPrime Full-Length cDNA Amplification Kit v1 (Lexogen GmbH), with the combination of the SMARTer PCR cDNA Synthesis kit; iii) TeloPrime Full-Length cDNA Amplification Kit v2 (Lexogen GmbH), with the combination of the SMARTer PCR cDNA Synthesis kit. The former protocol selectively synthesizes cDNA molecules from transcripts with a poly(A) tail and uses cDNA primers (5’ AAGCAGTGGTATCAACGCAGAGTACATGGGG; 3’ GTACTCTGCGTTGATACCACTGCTT). In contrast, the latter two protocols synthesize cDNA molecules from transcripts with both a 5’ CAP and a poly(A) tail, and uses cDNA primers (5’ TGGATTGATATGTAATACGACTCACTATAG; 3’ GTACTCTGCGTTGATACCACTGCTT). The two cDNA primer sets differ only in the 5’end, while the segments for the 3’end are the same. TeloPrime-based kit V2 includes updated reaction conditions in comparison with kit V1, while the method of cDNA generation remains unchanged. For each protocol, we used 1 ug homogeneous RNA as input to generate the cDNA library. The cDNAs were not size-selected, and PacBio libraries were prepared with the SMRTbell Template Prep Kit 1.0 (Pacific Biosciences) and sequenced on the PacBio Sequel I with Sequel DNA polymerase and binding kit 3.0 and sequencing chemistry version 3.0 for 1200 min. Each cDNA library was sequenced on one SMRT cell of the same sequencing run.

We analyzed the sub-reads for each SMRT cell separately following the IsoSeq3 pipeline (v3.4.0; https://github.com/PacificBiosciences/IsoSeq) and aligned the resulting isoforms to the mouse genome (GRCm39/mm39) using minimap2 (v2.24-r1122) (Li 2018), with the computational protocol indicated in the below text. The alignment results from the above three cDNA libraries were compared to the coordinates of transcripts annotated in Ensembl (v103) using gffcompare (v0.12.2) (Pertea and Pertea 2020).

Based on the detected isoforms in the Clontech library, we compare the transcripts with 5’ end truncation (*i.e.*, lacking at least the first exon, N=3,567) and the intact transcripts compared to reference annotation (N=4,199) at two different levels: i) the terminal exon length at 5’end; ii) the RNA structure property. The RNA structure property was measured as the minimum free energy (MFE) of the best predicted structure using RNAfold v2.6.4 with the following parameters “-p –d2 –noLP” (Gruber et al. 2008).

### PacBio Iso-Seq and Illumina RNA-Seq sequencing

The initial experimental tests showed that our newly developed TeloPrime-based cDNA library preparation protocol could provide a better solution to enrich for actual full-length transcripts, compared to the standard PacBio cDNA library preparation protocol. Hence, the TeloPrime Full-Length cDNA Amplification Kit v2 (Lexogen GmbH) was utilized to construct PacBio IsoSeq cDNA libraries. One µg total RNA from each individual was used as input, and double-strand cDNA was produced by following the manufacturer’s instructions, except that an alternative oligo-dT primer from the SMARTer PCR cDNA Synthesis kit (Clontech Laboratories, Inc.), which also included a random 10mer sequence as a unique molecule identifier (UMI) after each sequence. The cDNAs were not size-selected, and PacBio libraries were prepared with the SMRTbell Template Prep Kit 1.0 (Pacific Biosciences) and sequenced on the PacBio Sequel I with Sequel DNA polymerase and binding kit 3.0 and sequencing chemistry version 3.0 for 1200 min. Each library was sequenced on three SMRT cells to achieve sufficient coverage.

Poly(A) RNA from each sample was enriched from 1 µg total RNA by the NEBNext® Poly(A) mRNA Magnetic Isolation Module (Catalog #: E7490, New England Biolabs Inc.). RNA-Seq libraries were prepared using NEBNext Ultra™ II Directional RNA Library Prep Kit for Illumina (Catalog #: E7760, New England Biolabs Inc.), according to manufacturer’s instructions. A total of eleven PCR cycles were applied to enrich library concentration. Sequencing-by-synthesis was done at the HiSeq3000 system in paired-end mode 2 x 150bp. Raw sequencing outputs were converted to fastq files with bcl2fastq (v2.17.1.14).

### RNA-Seq read QC and data processing

We trimmed and filtered the low-quality raw fastq reads for each sample separately using the fastp program (v0.20.0; ––cut_front ––average_qual 20 ––length_required 50) (Chen et al. 2018) and only included the paired-end reads with a minimum length of 50bp and average quality score of 20 for further analysis.

The filtered fastq reads were aligned to mouse GRCm39/mm39 reference genome sequence with STAR aligner v2.7.0e (Dobin et al. 2013), taking the mouse gene annotation in Ensembl v103 (Howe et al. 2021) into account at the stage of building the genome index (––runMode genomeGenerate ––sjdbOverhang 149). The STAR mapping procedure was performed in two-pass mode, and some of the filtering parameters were tweaked (personal communication with STAR developer: see https://github.com/alexdobin/STAR/issues/1147; ––runMode alignReads –twopassMode Basic ––outFilterMismatchNmax 30 ––scoreDelOpen –1 ––scoreDelBase –1 ––scoreInsOpen –1 ––scoreInsBase –1 –– seedSearchStartLmax 25 ––winAnchorMultimapNmax 100), in order to compensate the sequence divergences of individuals from various populations and species (Zhang et al. 2021). With this optimized mapping pipeline, a similar alignment rate was reached for all the samples (Supplemental Data S1C). The alignment bam files were taken for further analysis.

### SNP variants calling and individual relatedness analysis

We followed the general GATK version 4 Best Practices to call genetic variants from Illumina RNA-seq data. We first sorted the above alignment bam data using samtools v1.9 (Li et al. 2009), and marked duplicates using PICARD v2.8.0 (http://broadinstitute.github.io/picard). Reads with N in the cigar were split into multiple supplementary alignments and hard clips mismatching overhangs using the SplitNCigarReads function in GATK v4.1.9. By using BaseRecalibrator and ApplyBQSR functions in GATK, we further recalibrated base quality scores with SNP variants that were called with the genomic sequencing dataset of the mice individuals from the same populations (Zhang et al. 2021) to get analysis-ready reads. Following, we called raw genetic variants for each individual using the HaplotypeCaller function in GATK and jointly genotyped genetic variants for all the individuals using the GenotypeGVCFs function. We only retained genetic variants that passed the hard filter “QD < 2.0 || FS > 60.0 || MQ < 40.0 || MQRankSum < –12.5 || ReadPosRankSum < –8.0 || SOR > 3.0” for further analysis (Zhang et al. 2021).

We used the relatedness2 option of VCFtools v0.1.14 (Danecek et al. 2011) to assess pairwise individual relatedness for all possible pairs of mice individuals, based on only bi-allelic SNP variants with no more than 20% missing data and locating within complete linear chromosomes, thinned to 1 SNP every 1 Mb (Harr et al. 2016). We defined that relatedness category based on the expected ranges of kinship coefficients (‘Phi’), according to the primary reference of the KING method (Manichaikul et al. 2010). The detailed relatedness information of all possible individual pairs can be found in Supplemental Data S1E.

### Iso-Seq read QC and data processing

We analyzed the raw sub-reads for each SMRT cell separately following the IsoSeq3 pipeline (v3.4.0). Circular consensus sequences (CCS) were generated from sub-reads using the CCS module in polish mode (v6.0.0; ––minPasses 3 ––minLength 50 ––maxLength 1000000 ––minPredictedAccuracy 0.99) of the IsoSeq3 pipeline, and the CCS reads generated in three SMRT cells for the same sample were merged. We trimmed the corresponding cDNA primers as shown above and orientated the CCS reads using the lima program with the specialized IsoSeq mode (v2.0.0; ––isoseq). The 10mer UMI following each CCS read was tagged and removed using the tag module of the IsoSeq3 pipeline (––design T-10U). Following this, we identified the processed CCS reads as full-length and non-chimeric (FLNC), based on the presence of a ploy(A) tail and absence of concatemer using the refine module of the IsoSeq3 pipeline (––require-polya). We further performed PCR deduplication based on the UMI tag information using the dedup module of the IsoSeq3 pipeline (default parameters). After this deduplication step, only one consensus FLNC sequence per founder molecule in the sample was kept. We then performed *de novo* clustering of the above reads using the cluster module of the IsoSeq3 pipeline and kept only the high-confidence isoforms supported by at least 2 FLNC reads for further analysis.

We aligned these isoforms of each sample to the GRCm39/mm39 reference genome sequence using the minimap2 (v2.24-r1122; –ax splice:hq –uf ––secondary=no –C5 –O6,24 –B4) (Li 2018), with parameter setting following the best practice of Cupcake pipeline (Gordon et al. 2015). The alignment bam files were further sorted using the samtools program (v1.9) (Li et al. 2009). We collapsed redundant isoform models for each sample based on the above sorted alignment coordinate information using the collapse module of the TAMA program (-d merge_dup –x capped –m 5 –a 1000 –z 30 –sj sj_priority) (Kuo et al. 2020). The rationales for defining redundant isoform models are shown in the Supplemental Text of Methods. Finally, isoforms across all 48 samples were merged into a single non-redundant transcriptome using the merge module of the TAMA program (-d merge_dup –m 5 –a 1000 –z 30) (Kuo et al. 2020), with the same parameter setting as the above collapse step.

### Iso-Seq transcriptome classification and filtering

We performed the quality control analysis for the above merged non-redundant PacBio Iso-Seq transcriptome using SQANTI3 v4.2 (Tardaguila et al. 2018), with the input datasets of Ensembl v103 (Howe et al. 2021) GRCm39/mm39 reference genome and gene annotation, FLNC read counts, Isoform expression levels and STAR output alignment bam files and splice junction files from RNA-Seq short reads, mouse transcription start sites (TSS) collected in refTSS database v3.1 (Abugessaisa et al. 2019), and curated set of poly(A) sites and poly(A) motifs in PolyASite portal v2.0 (Herrmann et al. 2020).

We filtered out isoforms of potential artifacts mainly by following (Tardaguila et al. 2018). Mono-exonic transcripts were excluded, as they tend more likely to be experimental or technical artifacts (Tardaguila et al. 2018). Isoforms with unreliable 3’end because of a possible intrapriming event (intrapriming rate above 0.6) were also removed from the dataset. We kept the remaining isoforms that met both of the following criteria: 1) no junction is not labeled as RT-Switching; 2) all junctions are either canonical (GT/AG; GC/AG; AT/AC) or supported by at least three spanning reads based on STAR junction output file. All isoforms that passed the above filters were taken for further analysis.

Given their matching status to the Ensembl mouse transcriptome v103 (Howe et al. 2021), the above isoforms were classified into eight distinct categories using SQANTI3 (Tardaguila et al. 2018): i) Full Splice Match (FSM, matching perfectly to a known isoform); ii) Incomplete Splice Match (ISM, matching to a subsection of a known isoform); iii) Novel In Category (NIC, with known splice sites but novel splice junctions); iv) Novel Not in Category (NNC, with at least one unannotated splice site); v) Fusion (F, fusion of adjacent transcripts); vi) Genic (G, overlapping with intron); vii) Antisense (A, on the antisense strand of an annotated gene); viii) Intergenic (I, within the intergenic region). The isoforms matching perfectly to a whole (FSM) or subsection (ISM) of reference annotated transcripts are designated known isoforms, and the others as novel isoforms.

We evaluated the reliability of isoforms from each category, based on the reference annotation and empirical information, from three distinct aspects separately: i) transcription start site (TSS); ii) transcript end site (TES); iii) splice junction (SJ). Compared to the well-support FSM isoforms, the isoforms from other categories show reduced confidence levels in terms of TSS (Supplemental Figure S5B) but no reduction for TES and SJ (Supplemental Figure S5D and 5E). This might hint at a failure to capture accurate TSS for those transcripts (Tardaguila et al. 2018). For instance, it is still possible that some ISM isoforms came from partial fragments due to the imperfect targeting of 5’ CAP or the degradation of transcripts in the later steps. To address this concern, we further excluded the non-FSM transcripts without support for TSSs (Supplemental Figure S5C).

### Rarefaction and subsampling

We investigated whether the PacBio Iso-Seq data provide sufficient coverage to detect all the isoform diversity for the given sequencing depth of a single individual and the number of sampled individuals in the experiment. Concerning the sequencing depth at the individual level, we randomly selected one sample from each of the seven assayed populations and subsampled portions of FLNC reads from each sample chosen for 100 times, ranging from 5% to 100%, at 5% intervals, and computed the fraction and variance of detected isoforms for each round of subsampling. Regarding the number of sampled individuals, we subsampled subsets of all the 48 assayed individuals for 100 times, ranging from 2 to 48, at an interval of 2, and computed the fraction and variance of detected isoforms for each round of subsampling.

We tested two alternative models to determine whether the number of detected isoforms would continue to increase or has approached saturation (Neme and Tautz 2016): a generalized linear model with logarithmic behavior (ever-increasing) or a self-starting nonlinear regression model (saturating). The best fit was decided based on the minimum BIC value between the two models, and the saturating model was the best fit for both lines of analysis. All the analyses were performed in R v4.2.3, using the functions glm, nls, SSasymp, and BI from the “stats” package (Team 2022).

### Test on the influence of noise on isoform diversity

We analyzed the influence of noise on isoform diversity using two different tests. For the first test, we extracted the list of isoforms supported by singleton FLNC reads in the de novo clustering step and filtered out potential artifacts using the same procedure as shown above. We performed the rarefaction analysis to test whether the number of singleton-supported isoforms has reached saturation with the sequencing depth at the individual level, following the abovementioned pipeline. We further compared the saturation curves between the isoforms represented by singleton FLNC reads and those represented by two or more FLNC reads (*i.e.*, high-confidence isoforms).

For the second test, we annotated the inserted transposon element (TE) fragments in the singleton and high-confidence isoforms using Repbase eukaryotes library v16.02 (Bao et al. 2015), which represents the largest TEs database for eukaryotes. We used Repeatmasker v4.0.5 (Tempel 2012) to mask the isoforms against the Repbase database with parameters “-nolow –no_is –norna –parallel 1”. Only the TE fragments longer than 10bp in length were included for further analysis. As several TE types might share the same coordinates, we removed overlapping TEs to retain a single most appropriate TE based on a series of criteria: i) if a TE insertion is entirely matching with another one with a different TEs type, the one with a larger mapping score was retained; ii) If two TE fragments overlapped partially, the overlapped part was assigned to the one with larger mapping score, then remaining part for the other TE, and was deleted if shorter than 10bp in length. This filtering was repeated across all pairs of overlapping domains until the defined conditions were satisfied. The TE insertion information for high-confidence and singleton isoforms can be found in Supplementary Data S2 and S3.

### Association between transcript pattern and population divergence

Based on three lines of analysis, we explored the association between individual isoform presence patterns and population divergence. Firstly, we calculated and compared the fraction of shared isoforms between all possible pairwise combinations from different species, from different subspecies but the same species (n = 448), from different populations but the same subspecies, and from the same populations. Secondly, we performed PCA on the individual isoform landscape using the R package ggfortify v0.4.16 (https://cran.r-project.org/web/packages/ggfortify/index.html). Additional PCAs were conducted for the subspecies *M. m. domesticus*, given that more closely related populations were assayed in this study.

Lastly, we built a phylogenetic tree for the seven assayed populations using the R package “ape” v5.7-1 (Paradis and Schliep 2019), based on each population’s presence/absence matrix of fixed isoforms. Euclidean distance was used as the distance measure between each pair of populations, and the neighbor-joining tree estimation function was used to build the phylogenetic relations. The boot.phylo function implemented in the same R package was used to perform 1,000 bootstrap replications, and population split nodes of high confidence were taken as the ones with at least 70% bootstrap support values. For comparison, we constructed another phylogenetic tree with the SNP variants that were called from the Illumina RNA-Seq dataset generated in this study (shown in the above text), for which the same set of mice individuals was used. To reduce computation complexity, we performed LD pruning on the SNP data set by using PLINK v1.90b4.6 (Purcell et al. 2007), removing one of a pair of SNPs with LD ≥0.2 in a sliding window of 500 SNPs and stepwise of 100 SNPs. The same procedure was followed to build the phylogenetic tree, as it was done for the analysis of isoforms.

### Detection of population differentiation isoform markers

We calculated population differentiation index (PDI) values for each isoform using a similar approach to analyze population differentiating retroCNVs (Zhang and Tautz 2022). A larger PDI value generally indicates greater power to use the presence status of the respective isoform to distinguish populations. The computation of the PDI value for each isoform follows:

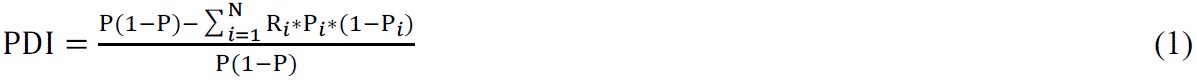

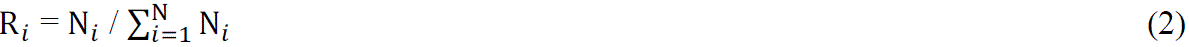

In the first equation, N represents the number of assayed populations (*i.e.*, 7), P*_i_* is the frequency of the focal isoform in the *i*th population, and P is the overall frequency in all the assayed populations. The definition of R*_i_* is shown in the second equation and is computed as the number of individuals of the *i*th population (N*_i_*) divided by the total number of individuals from all populations. We tested the significance of the PDI value for each isoform with a randomization test in which individual labels were randomly shuffled 1,000 times while population sizes remained the same. We calculated the p-value by comparing the observed PDI for each isoform against the PDI values from the randomized null distribution. Isoforms with significant PDI values after multiple testing corrections (FDR < 0.05) were considered population differentiation isoform markers.

### Feature characterization of all the isoforms

We characterized the features of all the detected isoforms from seven different aspects: i) exon number; ii) isoform length; iii) CDS length; iv) fraction of coding isoforms; v) fraction of NMD isoforms; vi) the number of individuals with expression; vii) isoform expression level. All these feature results were extracted from the output of SQANTI3 analysis.

In the SQANTI3 pipeline (Tardaguila et al. 2018), the potential coding capacity and ORFs from the isoform sequences were predicted using the GeneMarkS-T (GMST) algorithm (Tang et al. 2015). An NMD isoform is designated if there’s a predicted ORF, and the CDS ends at least 50bp before the last junction for the respective isoform. The expression level of each isoform for each sample was computed based on the number of supported FLNC reads and normalized in the unit of TPM (transcript per million). Isoforms with expression in respective tissues were defined as the ones with non-zero TPM values. In addition, we quantified the expression of each isoform using another program named Kallisto v0.46.2 (Bray et al. 2016), for which the expression quantification was based on the alignment of Illumina RNA-Seq data to the merged isoform dataset derived from PacBio Iso-Seq data, as shown in the above text.

### Feature characterization of gene loci with different number of isoforms

We classified all the detected gene loci into three roughly similar-sized classes, based on their splicing isoform numbers: i) Few-isoform Genes (FG, with no more than two isoforms, n = 5,372), ii) Many-isoform Genes (MG, with isoform number between three and eight, n = 5,012), and iii) Plenty-isoform Genes (PG, with no less than nine isoforms, n = 4,628). We characterized the features of the gene loci with different numbers of isoforms from three aspects, *i.e.*, gene expression pattern, gene age distribution pattern, and gene ontology (GO) functional term enrichment.

Firstly, we exploited two Illumina RNA-seq datasets for the gene expression pattern analysis: i) the one generated in this study for the whole brain of 48 mice individuals; ii) a published dataset from thirteen tissues (adipose tissue, adrenal gland, brain, heart, kidney, liver, lung, pancreas, sigmoid colon, small intestine, spleen, ovary, and testis) of the *M. m. domesticus* C57BL/6 inbred line (Lin et al. 2014). The raw fastq data for both datasets were trimmed, filtered, and aligned to the mouse GRCm39/mm39 reference genome, as shown in the above text. We counted the fragments uniquely mapped to the annotated genes in Ensembl v103 using featureCounts v1.6.3 (Liao et al. 2014) in reversely stranded mode, excluding multi-mapping reads and those with alignment quality below 5 (-s 2 –Q 5). Gene expression levels were calculated and transformed into a log2 scale of FPKM (Fragments Per Kilobase of transcript per Million mapped fragments) based on the gene’s isoform length and the number of mapped fragments in the data set from each sample. For the first dataset, the average gene expression level was computed as the mean value in all the assayed 48 samples. For the second dataset, the number of tissues with expression for each gene was defined as the count of tissues with non-zero FPKM values. The statistical significances of the expression patterns between pairwise groups of gene sets were computed using two-sided Wilcoxon rank sum tests.

Secondly, we analyzed and compared the evolutionary gene age pattern among three gene groups, taking advantage of the mouse gene age assignment data from Neme *et al*. (Neme and Tautz 2013). In brief, each mouse protein-coding gene was dated and given phylostratum (PS) assignment by inferring the absence and presence of orthologs along the phylogenetic tree. The statistical significances of PS values between pairwise groups of gene set were computed using two-sided Wilcoxon rank sum tests.

Lastly, we performed GO enrichment analysis on the genes from the PG group using GOSeq v1.50.0 (Young et al. 2010), without correction of gene length differences. Only significant BP (biological process) terms after multiple testing corrections (FDR < 0.05) were listed in this study.

### Calculation of isoform expression for highly versus lowly expressed genes

We focused on the genes with at least two isoforms in at least one individual of the *M. musculus* populations, resulting in a total of 9,111 gene loci for further analysis. The isoform data in outgroups were excluded for this since many top-expressed isoforms can change in these outgroups. We used the normalized FLNC read counts (TPM) as a proxy for the expression level of isoforms and the sum of expression of all isoforms belonging to a given gene locus. The final isoform/gene expression values were averaged across all the *M. musculus* individuals.

We calculated each gene’s top expressed isoform read counts (T) by the read counts of the remaining isoforms (R) from the same genes. These calculations were performed for all the genes, the top 10% expressed genes and lowest 10% expressed genes, separately. The statistical significance on the ratio difference between the top 10% expressed genes and lowest 10% expressed genes was computed using Wilcoxon rank sum test.

### Analysis of local AS events

We used the SUPPA2 program (Trincado et al. 2018) to identify local alternative splicing (AS) events in the transcriptome. These local AS events are categorized into seven groups, including skipped exon (SE), retained intron (RI), alternative 5’ splice site (A5), alternative 3’ splice site (A3), mutually exclusive exon (MX), alternative first exon (AFE), and alternative last exon (ALE).

For the isoforms with predicted ORFs, we analyzed how the local AS events change the ORF structures. In case the start codon position of the ORF lands in the region of a local AS event, the focal AS event is defined to cause a change in the respective coding sequence. On the other hand, in case the start codon position of the ORF falls downstream of the local AS event, it does not influence the ORF structure.

## Data access

The raw PacBio Iso-Seq data and Illumina RNA-Seq data generated in this study have been submitted to the European Nucleotide Archive (ENA; https://www.ebi.ac.uk/ena) under study accession number PRJEB54001 (test experiment), PRJEB54000 and PRJEB53988 (the main experiment). Alignment bam files, GTF track data, and SNP VCF files are stored at the ftp site: https://www.user.gwdg.de/~evolbio/evolgen/wildmouse/mouse_population_isoform/ and supplemental data at https://edmond.mpg.de/dataset.xhtml?persistentId=doi:10.17617/3.3RLIHR All the essential computing codes are provided as Supplemental Data 8, and related data files are available at GitLab: https://gitlab.gwdg.de/wenyu.zhang/mouse_population_isoform/.

## Competing interest statement

The authors declare that they have no conflicts of interests.

## Supporting information

Combined supplemental files

## Acknowledgements

We appreciate Christine Pfeifle, Heike Harre, Milan Jovicic, and Mustafa Al-Ameer for mice breeding and handling. We thank Carsten Fortmann-Grote for computing assistance and Chen Xie for helpful discussion. We thank Henrik Kaessmann for helpful suggestions on the manuscript. Computing was supported by the high-performance computing clusters of the Max Planck Institute for Evolutionary Biology and School of Ecology and Environment at Northwestern Polytechnical University. This work was supported by institutional funding through the Max Planck Society to D.T. and W.Z., and the Fundamental Research Funds for the Central Universities in China (grant number: G2022KY05106), National Natural Science Foundation of China (grant number: 32370665), Guangdong Basic and Applied Basic Research Foundation (grant number: 2024A1515030117), and the Innovation Capability Support Plan Project in Shaanxi Province (grant number: 2024ZC-KJXX-038) to W.Z.

## Notes

https://edmond.mpg.de/dataset.xhtml?persistentId=doi:10.17617/3.3RLIHR

### Competing Interest Statement

The authors have declared no competing interest.

### Summary of Updates

The manuscript was substantially revised, including several additional analyses, which involved also additional scientists, who are now listed as further co-authors.

## References

1. Abugessaisa I, Noguchi S, Hasegawa A, Kondo A, Kawaji H, Carninci P, Kasukawa T. 2019. refTSS: A Reference Data Set for Human and Mouse Transcription Start Sites. J Mol Biol 431: 2407–2422.

2. Andersson R, Sandelin A. 2020. Determinants of enhancer and promoter activities of regulatory elements. Nat Rev Genet 21: 71–87.

3. Bao W, Kojima KK, Kohany O. 2015. Repbase Update, a database of repetitive elements in eukaryotic genomes. Mob DNA 6: 11.

4. Barbosa-Morais NL, Irimia M, Pan Q, Xiong HY, Gueroussov S, Lee LJ, Slobodeniuc V, Kutter C, Watt S, Colak R et al. 2012. The evolutionary landscape of alternative splicing in vertebrate species. Science 338: 1587–1593.

5. Bekpen C, Xie C, Tautz D. 2018. Dealing with the adaptive immune system during de novo evolution of genes from intergenic sequences. BMC Evol Biol 18: 121.

6. Benitiere F, Necsulea A, Duret L. 2024. Random genetic drift sets an upper limit on mRNA splicing accuracy in metazoans. Elife 13.

7. Bray NL, Pimentel H, Melsted P, Pachter L. 2016. Near-optimal probabilistic RNA-seq quantification. Nat Biotechnol 34: 525–527.

8. Byrne A, Cole C, Volden R, Vollmers C. 2019. Realizing the potential of full-length transcriptome sequencing. Philos Trans R Soc Lond B Biol Sci 374: 20190097.

9. Cartolano M, Huettel B, Hartwig B, Reinhardt R, Schneeberger K. 2016. cDNA Library Enrichment of Full Length Transcripts for SMRT Long Read Sequencing. PLoS One 11: e0157779.

10. Chang YF, Imam JS, Wilkinson MF. 2007. The nonsense-mediated decay RNA surveillance pathway. Annu Rev Biochem 76: 51–74.

11. Chen S, Zhou Y, Chen Y, Gu J. 2018. fastp: an ultra-fast all-in-one FASTQ preprocessor. Bioinformatics 34: i884–i890.

12. Danecek P, Auton A, Abecasis G, Albers CA, Banks E, DePristo MA, Handsaker RE, Lunter G, Marth GT, Sherry ST et al. 2011. The variant call format and VCFtools. Bioinformatics 27: 2156–2158.

13. Ding C, Yan X, Xu M, Zhou R, Zhao Y, Zhang D, Huang Z, Pan Z, Xiao P, Li H et al. 2022. Short-read and long-read full-length transcriptome of mouse neural stem cells across neurodevelopmental stages. Sci Data 9: 69.

14. Dobin A, Davis CA, Schlesinger F, Drenkow J, Zaleski C, Jha S, Batut P, Chaisson M, Gingeras TR. 2013. STAR: ultrafast universal RNA-seq aligner. Bioinformatics 29: 15–21.

15. Eisenberg E, Levanon EY. 2003. Human housekeeping genes are compact. Trends Genet 19: 362–365.

16. Ferrandez-Peral L, Zhan X, Alvarez-Estape M, Chiva C, Esteller-Cucala P, Garcia-Perez R, Julia E, Lizano E, Fornas O, Sabido E et al. 2022. Transcriptome innovations in primates revealed by single-molecule long-read sequencing. Genome Res doi:10.1101/gr.276395.121.

17. Gordon SP, Tseng E, Salamov A, Zhang J, Meng X, Zhao Z, Kang D, Underwood J, Grigoriev IV, Figueroa M et al. 2015. Widespread Polycistronic Transcripts in Fungi Revealed by Single-Molecule mRNA Sequencing. PLoS One 10: e0132628.

18. Grau-Bove X, Ruiz-Trillo I, Irimia M. 2018. Origin of exon skipping-rich transcriptomes in animals driven by evolution of gene architecture. Genome Biol 19: 135.

19. Gruber AR, Lorenz R, Bernhart SH, Neubock R, Hofacker IL. 2008. The Vienna RNA websuite. Nucleic Acids Res 36: W70–74.

20. Guenet JL, Bonhomme F. 2003. Wild mice: an ever-increasing contribution to a popular mammalian model. Trends Genet 19: 24–31.

21. Gupta I, Collier PG, Haase B, Mahfouz A, Joglekar A, Floyd T, Koopmans F, Barres B, Smit AB, Sloan SA et al. 2018. Single-cell isoform RNA sequencing characterizes isoforms in thousands of cerebellar cells. Nat Biotechnol doi:10.1038/nbt.4259.

22. Harr B, Karakoc E, Neme R, Teschke M, Pfeifle C, Pezer Z, Babiker H, Linnenbrink M, Montero I, Scavetta R et al. 2016. Genomic resources for wild populations of the house mouse, Mus musculus and its close relative Mus spretus. Sci Data 3: 160075.

23. Harr B, Turner LM. 2010. Genome-wide analysis of alternative splicing evolution among Mus subspecies. Mol Ecol 19 **Suppl 1**: 228–239.

24. Herrmann CJ, Schmidt R, Kanitz A, Artimo P, Gruber AJ, Zavolan M. 2020. PolyASite 2.0: a consolidated atlas of polyadenylation sites from 3’ end sequencing. Nucleic Acids Res 48: D174–D179.

25. Howe KL, Achuthan P, Allen J, Allen J, Alvarez-Jarreta J, Amode MR, Armean IM, Azov AG, Bennett R, Bhai J et al. 2021. Ensembl 2021. Nucleic Acids Res 49: D884–D891.

26. Huang KK, Huang J, Wu JKL, Lee M, Tay ST, Kumar V, Ramnarayanan K, Padmanabhan N, Xu C, Tan ALK et al. 2021. Long-read transcriptome sequencing reveals abundant promoter diversity in distinct molecular subtypes of gastric cancer. Genome Biol 22: 44.

27. Ilik IA, Glazar P, Tse K, Brandl B, Meierhofer D, Muller FJ, Smith ZD, Aktas T. 2024. Autonomous transposons tune their sequences to ensure somatic suppression. Nature doi:10.1038/s41586-024-07081-0.

28. Joglekar A, Prjibelski A, Mahfouz A, Collier P, Lin S, Schlusche AK, Marrocco J, Williams SR, Haase B, Hayes A et al. 2021. A spatially resolved brain region– and cell type-specific isoform atlas of the postnatal mouse brain. Nat Commun 12: 463.

29. Keren H, Lev-Maor G, Ast G. 2010. Alternative splicing and evolution: diversification, exon definition and function. Nat Rev Genet 11: 345–355.

30. Kuo RI, Cheng Y, Zhang R, Brown JWS, Smith J, Archibald AL, Burt DW. 2020. Illuminating the dark side of the human transcriptome with long read transcript sequencing. BMC Genomics 21: 751.

31. Kuo RI, Tseng E, Eory L, Paton IR, Archibald AL, Burt DW. 2017. Normalized long read RNA sequencing in chicken reveals transcriptome complexity similar to human. BMC Genomics 18: 323.

32. Landry JR, Mager DL, Wilhelm BT. 2003. Complex controls: the role of alternative promoters in mammalian genomes. Trends Genet 19: 640–648.

33. Lebrigand K, Magnone V, Barbry P, Waldmann R. 2020. High throughput error corrected Nanopore single cell transcriptome sequencing. Nat Commun 11: 4025.

34. Leong AZ, Lee PY, Mohtar MA, Syafruddin SE, Pung YF, Low TY. 2022. Short open reading frames (sORFs) and microproteins: an update on their identification and validation measures. J Biomed Sci 29: 19.

35. Leung SK, Jeffries AR, Castanho I, Jordan BT, Moore K, Davies JP, Dempster EL, Bray NJ, O’Neill P, Tseng E et al. 2021. Full-length transcript sequencing of human and mouse cerebral cortex identifies widespread isoform diversity and alternative splicing. Cell Rep 37: 110022.

36. Li H. 2018. Minimap2: pairwise alignment for nucleotide sequences. Bioinformatics 34: 3094–3100.

37. Li H, Handsaker B, Wysoker A, Fennell T, Ruan J, Homer N, Marth G, Abecasis G, Durbin R, Genome Project Data Processing S. 2009. The Sequence Alignment/Map format and SAMtools. Bioinformatics 25: 2078–2079.

38. Liao Y, Smyth GK, Shi W. 2014. featureCounts: an efficient general purpose program for assigning sequence reads to genomic features. Bioinformatics 30: 923–930.

39. Lin L, Shen S, Jiang P, Sato S, Davidson BL, Xing Y. 2010. Evolution of alternative splicing in primate brain transcriptomes. Hum Mol Genet 19: 2958–2973.

40. Lin S, Lin Y, Nery JR, Urich MA, Breschi A, Davis CA, Dobin A, Zaleski C, Beer MA, Chapman WC et al. 2014. Comparison of the transcriptional landscapes between human and mouse tissues. Proc Natl Acad Sci U S A 111: 17224–17229.

41. Ling Z, Brockmoller T, Baldwin IT, Xu S. 2019. Evolution of Alternative Splicing in Eudicots. Front Plant Sci 10: 707.

42. Malko DB, Makeev VJ, Mironov AA, Gelfand MS. 2006. Evolution of exon-intron structure and alternative splicing in fruit flies and malarial mosquito genomes. Genome Res 16: 505–509.

43. Manichaikul A, Mychaleckyj JC, Rich SS, Daly K, Sale M, Chen WM. 2010. Robust relationship inference in genome-wide association studies. Bioinformatics 26: 2867–2873.

44. Mudge JM, Frankish A, Fernandez-Banet J, Alioto T, Derrien T, Howald C, Reymond A, Guigo R, Hubbard T, Harrow J. 2011. The origins, evolution, and functional potential of alternative splicing in vertebrates. Mol Biol Evol 28: 2949–2959.

45. Murat F, Mbengue N, Winge SB, Trefzer T, Leushkin E, Sepp M, Cardoso-Moreira M, Schmidt J, Schneider C, Mossinger K et al. 2023. The molecular evolution of spermatogenesis across mammals. Nature 613: 308–316.

46. Naftaly AS, Pau S, White MA. 2021. Long-read RNA sequencing reveals widespread sex-specific alternative splicing in threespine stickleback fish. Genome Res 31: 1486–1497.

47. Neme R, Tautz D. 2013. Phylogenetic patterns of emergence of new genes support a model of frequent de novo evolution. BMC Genomics 14: 117.

48. Neme R, Tautz D. 2016. Fast turnover of genome transcription across evolutionary time exposes entire non-coding DNA to de novo gene emergence. Elife 5.

49. Nilsen TW, Graveley BR. 2010. Expansion of the eukaryotic proteome by alternative splicing. Nature 463: 457–463.

50. Parada GE, Munita R, Cerda CA, Gysling K. 2014. A comprehensive survey of non-canonical splice sites in the human transcriptome. Nucleic Acids Res 42: 10564–10578.

51. Paradis E, Schliep K. 2019. ape 5.0: an environment for modern phylogenetics and evolutionary analyses in R. Bioinformatics 35: 526–528.

52. Pertea G, Pertea M. 2020. GFF Utilities: GffRead and GffCompare. F1000Res 9.

53. Pezer Z, Harr B, Teschke M, Babiker H, Tautz D. 2015. Divergence patterns of genic copy number variation in natural populations of the house mouse (Mus musculus domesticus) reveal three conserved genes with major population-specific expansions. Genome Res 25: 1114–1124.

54. Phifer-Rixey M, Nachman MW. 2015. Insights into mammalian biology from the wild house mouse Mus musculus. Elife 4.

55. Pickrell JK, Pai AA, Gilad Y, Pritchard JK. 2010. Noisy splicing drives mRNA isoform diversity in human cells. PLoS Genet 6: e1001236.

56. Purcell S, Neale B, Todd-Brown K, Thomas L, Ferreira MA, Bender D, Maller J, Sklar P, de Bakker PI, Daly MJ et al. 2007. PLINK: a tool set for whole-genome association and population-based linkage analyses. Am J Hum Genet 81: 559–575.

57. Rogers TF, Palmer DH, Wright AE. 2021. Sex-Specific Selection Drives the Evolution of Alternative Splicing in Birds. Mol Biol Evol 38: 519–530.

58. Saudemont B, Popa A, Parmley JL, Rocher V, Blugeon C, Necsulea A, Meyer E, Duret L. 2017. The fitness cost of mis-splicing is the main determinant of alternative splicing patterns. Genome Biol 18: 208.

59. Sharon D, Tilgner H, Grubert F, Snyder M. 2013. A single-molecule long-read survey of the human transcriptome. Nat Biotechnol 31: 1009–1014.

60. Staubach F, Lorenc A, Messer PW, Tang K, Petrov DA, Tautz D. 2012. Genome patterns of selection and introgression of haplotypes in natural populations of the house mouse (Mus musculus). PLoS Genet 8: e1002891.

61. Tang S, Lomsadze A, Borodovsky M. 2015. Identification of protein coding regions in RNA transcripts. Nucleic Acids Res 43: e78.

62. Tardaguila M, de la Fuente L, Marti C, Pereira C, Pardo-Palacios FJ, Del Risco H, Ferrell M, Mellado M, Macchietto M, Verheggen K et al. 2018. SQANTI: extensive characterization of long-read transcript sequences for quality control in full-length transcriptome identification and quantification. Genome Res doi:10.1101/gr.222976.117.

63. Team RC. 2022. R: A language and environment for statistical computing. R Foundation for Statistical Computing, Vienna, Austria.

64. Tempel S. 2012. Using and understanding RepeatMasker. Methods Mol Biol 859: 29–51.

65. Tian B, Manley JL. 2017. Alternative polyadenylation of mRNA precursors. Nat Rev Mol Cell Biol 18: 18–30.

66. Trincado JL, Entizne JC, Hysenaj G, Singh B, Skalic M, Elliott DJ, Eyras E. 2018. SUPPA2: fast, accurate, and uncertainty-aware differential splicing analysis across multiple conditions. Genome Biol 19: 40.

67. Verta JP, Jacobs A. 2022. The role of alternative splicing in adaptation and evolution. Trends Ecol Evol 37: 299–308.

68. Wan Y, Larson DR. 2018. Splicing heterogeneity: separating signal from noise. Genome Biol 19: 86.

69. Young MD, Wakefield MJ, Smyth GK, Oshlack A. 2010. Gene ontology analysis for RNA-seq: accounting for selection bias. Genome Biol 11: R14.

70. Zhang W, Landback P, Gschwend AR, Shen B, Long M. 2015. New genes drive the evolution of gene interaction networks in the human and mouse genomes. Genome Biol 16: 202.

71. Zhang W, Tautz D. 2022. Tracing the Origin and Evolutionary Fate of Recent Gene Retrocopies in Natural Populations of the House Mouse. Mol Biol Evol 39.

72. Zhang W, Xie C, Ullrich K, Zhang YE, Tautz D. 2021. The mutational load in natural populations is significantly affected by high primary rates of retroposition. Proc Natl Acad Sci U S A 118.

