## Supplementary material for "Full-length RNA transcript sequencing traces brain isoform diversity in house mouse natural populations": Combined supplemental files

**Supplemental Text of Methods**

**Supplemental Figures and Tables**

- **Figure S1** Comparison of intact transcripts and the degraded transcripts from 5'end
- **Figure S2** Schematic pipeline of the PacBio Iso-Seq and Illumina RNA-Seq data analysis
- **Figure S3** Schematic diagram for the collapsing of redundant transcript models
- **Figure S4** Calculation of coverage ratios of TSSs
- **Figure S5** Support level for transcripts from different structural categories
- **Figure S6** Features of all Iso-Seq detected transcripts compared to reference annotated transcripts
- **Figure S7** Support level of transcripts from different populations
- **Figure S8** Rarefaction analysis for singletons from individuals of the different populations
- **Figure S9** Rarefaction analysis for high confidence isoforms from individuals of the different populations
- **Figure S10** Comparison of TE insertions between singleton isoforms and high-confidence isoforms
- **Figure S11** Population differentiation pattern based on higher-frequency isoforms
- **Figure S12** Distribution of structural categories of house mouse isoforms
- **Figure S13** Distribution of isoform and ORF numbers across all the seven assayed populations
- **Figure S14** Genomic features of gene loci with different numbers of isoforms
- **Table S1** Summary of sequencing statistics generated in this study
- **Table S2** Summary of statistics for the comparison analysis between known and novel transcripts

**SI References**

**Other supplemental materials for this manuscript include the following:**

- **Data S1:** Summary of the sampled mice individuals and sequencing datasets in this study

    Data S1A: Summary of sequencing statistics for the optimization experiment

    Data S1B: Individual information and sequencing statistics of PacBio Iso-Seq datasets

    Data S1C: Sequencing statistics of Illumina RNA-Seq datasets

    Data S1D: Publicly available transcriptomic sequencing data used in this study

    Data S1E: Pairwise individual relatedness estimates for all possible pairs of individuals

- **Data S2** The list of high-confidence isoforms detected in this study
- **Data S3** The list of singleton isoforms detected in this study
- **Data S4** Statistics on the known and novel isoforms for each mouse individual
- **Data S5** GO functional enrichment results of heavily spliced genes
- **Data S6** Ratio analysis for normalized isoform read counts (TPM)
- **Data S7** GTF data for isoforms conserved between house mouse and outgroups
- **Data S8** Essential computing code used in this study

### Supplemental Text of Methods

*Optimizing parameters to collapse redundant transcript models*

We utilized the "collapse" module of TAMA program (Kuo et al. 2020) to collapse redundant transcript models based on the alignment coordinate information (Supplemental Figure S3). The selection of the threshold of the amount of tolerance at the 5' end of the transcript for the grouping was based on the mean distance (roughly 2,000nt, thus choosing "–a 1000" given that collapsing process occurs in two directions) between adjacent transcription start sites (TSSs) of protein-coding genes detected from cap analysis gene expression sequencing (CAGE-Seq) data in mouse tissues (Xu et al. 2019). We chose the threshold of the amount of tolerance at the 3' end of the transcript for grouping (-z 30) by following (Xu and Zhang 2018), for which it was determined on the basis of the empirical evidence from PolyA-seq dataset (Derti et al. 2012).

Unlike the collapse of the flanking regions' terminal exons, the alignment error is the primary consideration in choosing the tolerance threshold at splice junction sites (Kuo et al. 2020). To determine this threshold, we retrieved all the cDNA sequences (with ≥2 exons) based on the mouse gene annotation in Ensembl v103 (Howe et al. 2021). We further introduced 1% of random single-site mutations in these sequences to mimic the sequenced Iso-Seq long reads, in accordance with the predicting accuracy (99%) of CCS reads documented in the IsoSeq3 pipeline. These simulated reads were aligned to GRCm39/mm39 reference genome sequence using minimap2 with the same parameters as shown in the main text. We calculated the distances between the aligned splice junction sites and the original annotation in Ensembl v103 (Supplemental Figure S3D). Most of the splice junctions (99.79%) were accurately aligned to the GRCm39/mm39 reference genome sequences, and almost none of them were aligned to distinct positions of 10bp further than the original annotations. Consequently, we chose "–m 5" as the cut-off to group transcripts with similar splice junction sites, given that collapsing process occurs in two directions.

*Selection of coverage ratio threshold to define "reliable" TSSs*

The coverage ratio inside and outside of an isoform should be greater than 1 (Supplemental Figure S4A). Consequently, this type of coverage ratio (Ratio_TSS) could possibly act as an index to evaluate the reliability of TSSs (Tardaguila et al. 2018). To test this hypothesis, we first generated a dataset of "pseudo" TSSs by randomly selecting 1,000 sites from exon regions annotated in GRCm39/mm39 reference based on Ensembl v103 (Howe et al. 2021). We measured the Ratio_TSS value as the mean coverage of the 100bp upstream and downstream of these artificial TSSs based on Illumina RNA-seq data, and took the maximum value of the ratios across all 48 samples (Tardaguila et al. 2018). Most of the Ratio_TSS values are close to 1, hinting that the majority of these sites are actually "fake" TSSs (Supplemental Figure S4B). As expected, the Ratio_TSS values of TSSs from reference annotation and filtered Iso-Seq isoforms detected in our study are significantly greater than those of "pseudo" TSSs (Supplemental Figure S4B), as most of these sites may represent "true" TSSs. In this study, we chose the median Ratio_TSS value of reference annotated TSSs (*i.e.*, 1.5) as the cutoff to evaluate the reliability of detected TSSs for the filtered Iso-Seq isoforms (Supplemental Figure S4B).

*Evaluation of the reliability of all the detected Iso-Seq isoforms*

We evaluated the reliability of Iso-Seq isoforms on the basis of reference annotation and empirical information, from three distinct aspects separately (Supplemental Figure S5A): i) Transcription start site (TSS); ii) Transcript End Site (TES); iii) splice junction (SJ). A given TSS is considered as reliable if any of the following criteria is met: i) with a distance of no more than 50bp to annotated TSS of any transcripts of the matched gene in GRCm39/mm39 reference genome from Ensembl v103 (Howe et al. 2021); ii) within any CAGE peak of mouse strains collected in refTSS database (v3.1) (Abugessaisa et al. 2019); iii) with a minimum coverage ratio 1.5 inside isoform and outside isoform (Tardaguila et al. 2018) (see above). Similarly, a reliable TES is defined as long as any of the following criteria is met: i) with a distance of no more than 50bp to annotated TES of any transcripts of the matched gene in GRCm39/mm39 reference genome from Ensembl v103 (Howe et al. 2021); ii) within any poly(A) peak of mouse strains collected in PolyASite portal (v2.0) (Herrmann et al. 2020); iii) with poly(A) motif in PolyASite portal (v2.0) (Herrmann et al. 2020) detectable within 50bp upstream of the given TES. We also assessed the quality of SJs by checking whether they are either canonical splicing signals (GT-AG, GC-AG, and AT-AC) or present in the GRCm39/mm39 reference genome annotation from Ensembl v103 (Howe et al. 2021).

Further, we compared the detected transcripts with those annotated in the GRCm39/mm39 reference genome from Ensembl v103 (Howe et al. 2021), built largely based on Illumina short-read RNA-Seq data. In comparison to the reference annotated transcripts, the transcripts derived from the PacBio Iso-Seq approach have a significantly larger number of exons (Supplemental Figure S6A, median 7 VS 5, two-sided Wilcoxon rank sum test, p-value $< 2.2 \times 10^{-16}$), and longer transcript lengths (Supplemental Figure S6B, median 1,887 VS 1,112, two-sided Wilcoxon rank sum test, p-value $< 2.2 \times 10^{-16}$). Nevertheless, the coding sequence (CDS) lengths from the coding transcripts of these two groups are more comparable (Supplemental Figure S6C, median value 870 VS 888, two-sided Wilcoxon rank sum test, p-value = 0.46). These findings further confirm the performance of PacBio Iso-Seq technology to better capture full-length transcripts (Kuo et al. 2020), especially for the regulatory UTR regions (Supplemental Figure S6D).

*Evaluation of the quality of detected isoforms across all the seven assayed populations*

We analyzed the features of detected isoforms from different populations, and found that a similar number of transcripts (mean = 21,867, corresponding to 8,209 genes) per sample can be identified for all the seven assayed natural populations (Table 1). Notably, approximately 0.8% (SD: 0.002) of detected gene loci and 25.5% (SD: 0.02) of detected transcripts from each sample in all seven populations were not represented in the GRCm39/mm39 Ensembl annotation (Table 1) (Howe et al. 2021). Among the detected transcripts per individual from each population, a consistent 92.5% (SD: 0.01) of them were predicted with coding capability defined by SQANTI3 (Tardaguila et al. 2018) (Table 1), corresponding to a similar value of 93.4% detected in the cortex from a lab mouse strain (Leung et al. 2021). We also found that around 5.5% (SD: 0.005) of these predicted coding transcripts bear a signal of nonsense-mediated decay (NMD), *i.e.*, with a premature stop codon. All these transcripts can be confirmed to a comparable large extent (>96%) by the experimental and empirical data for TSSs, TESs, and SJs, respectively (Supplemental Figure S7).

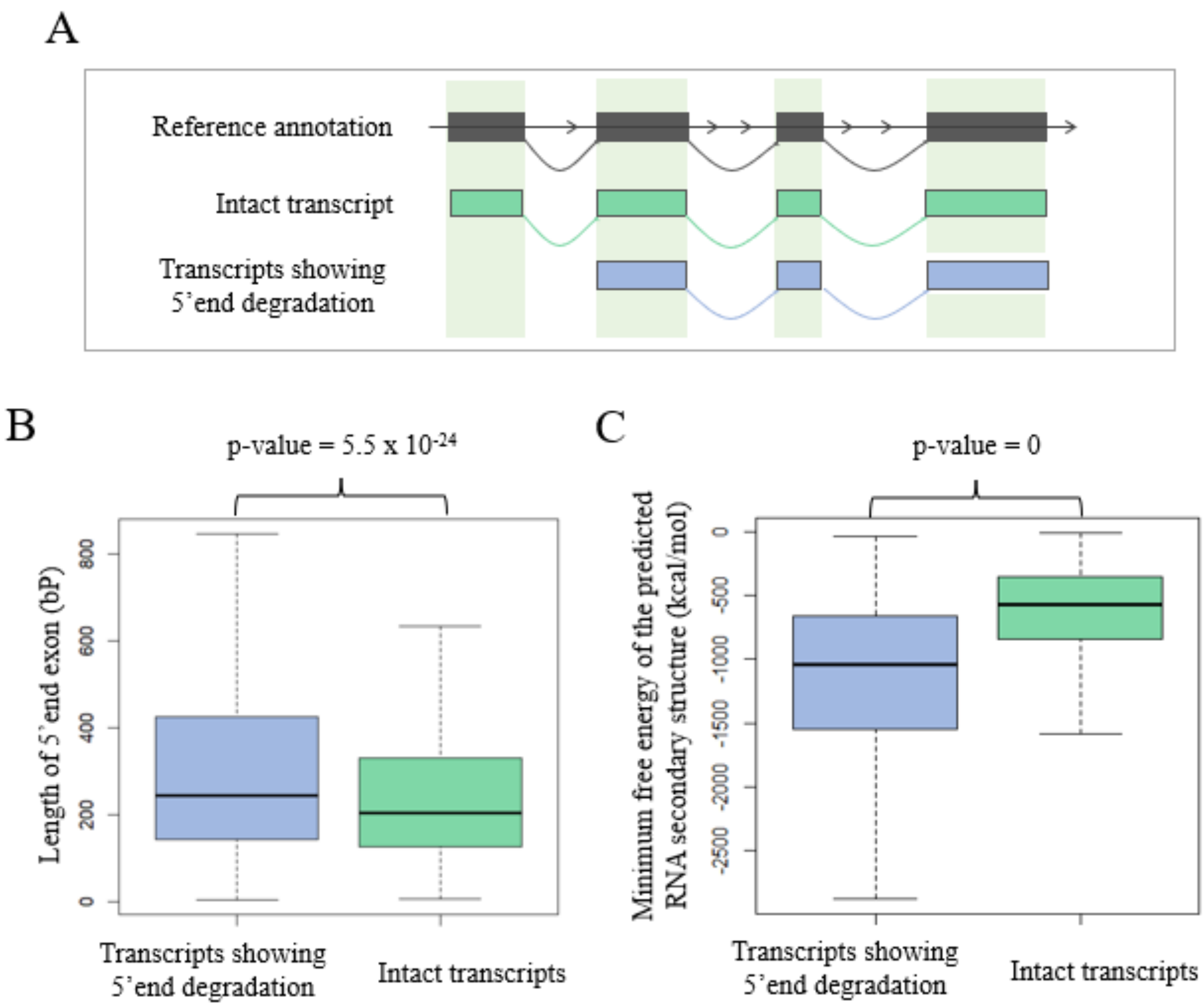


**Figure S1 Comparison of intact transcripts and the degraded transcripts from 5'end.** This figure shows the feature comparison between the reference transcripts having isoforms with 5'end exon degradation and having isoforms perfectly match to the reference annotation. Note that only the isoforms detected in the Clontech library preparation protocol were taken for analysis. (A) shows the schematic diagram of intact transcripts and transcript showing 5'end degradation. (B) and (C) show the comparison of 5' terminal exon length and minimum free energy of predicted RNA secondary structure of the two types of transcripts.

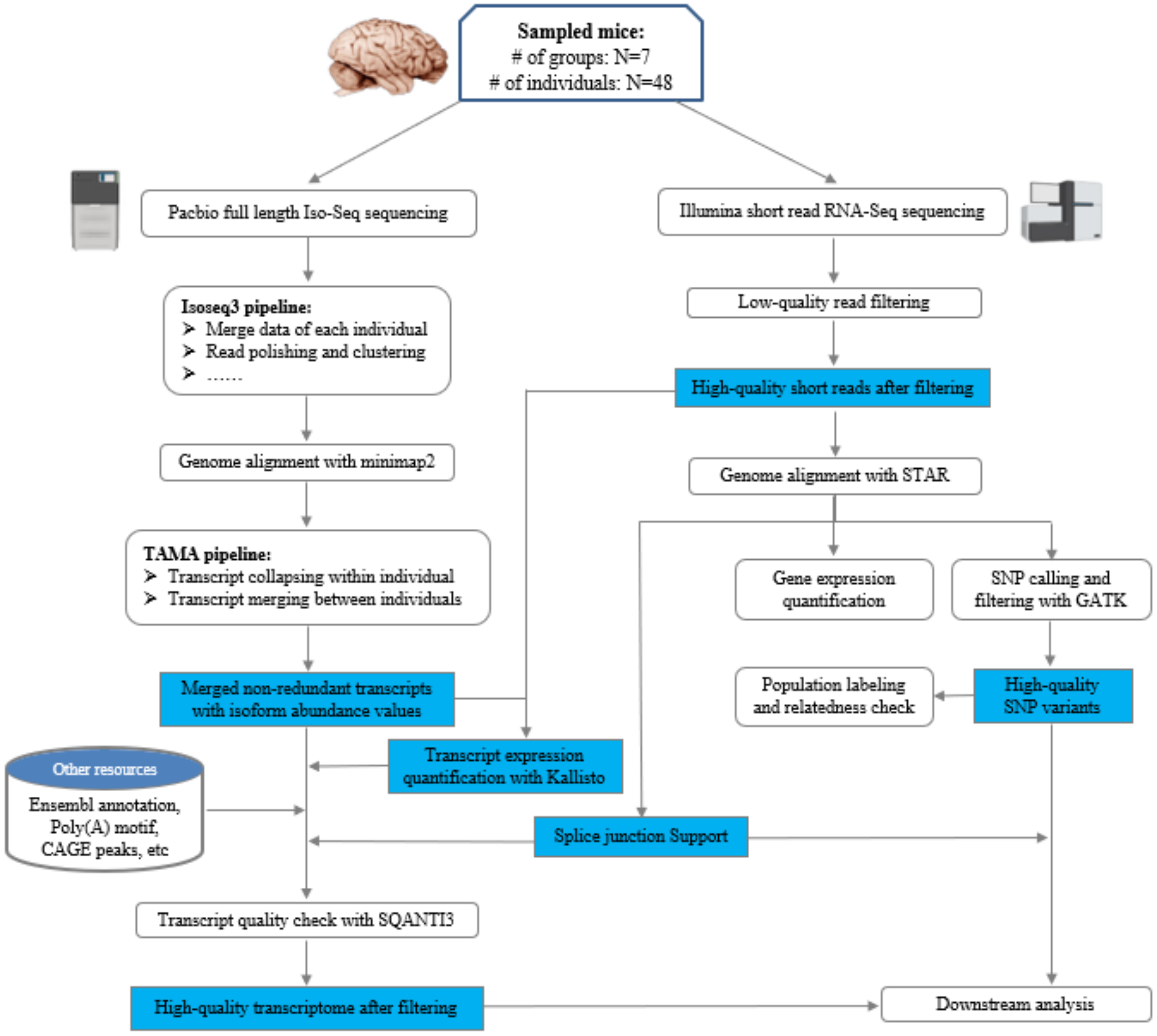


**Figure S2 Schematic pipeline of the PacBio Iso-Seq and Illumina RNA-Seq data analysis.** This pipeline mainly shows the steps to generate high-quality merged transcriptomes with the combination of Iso-Seq and RNA-seq data.

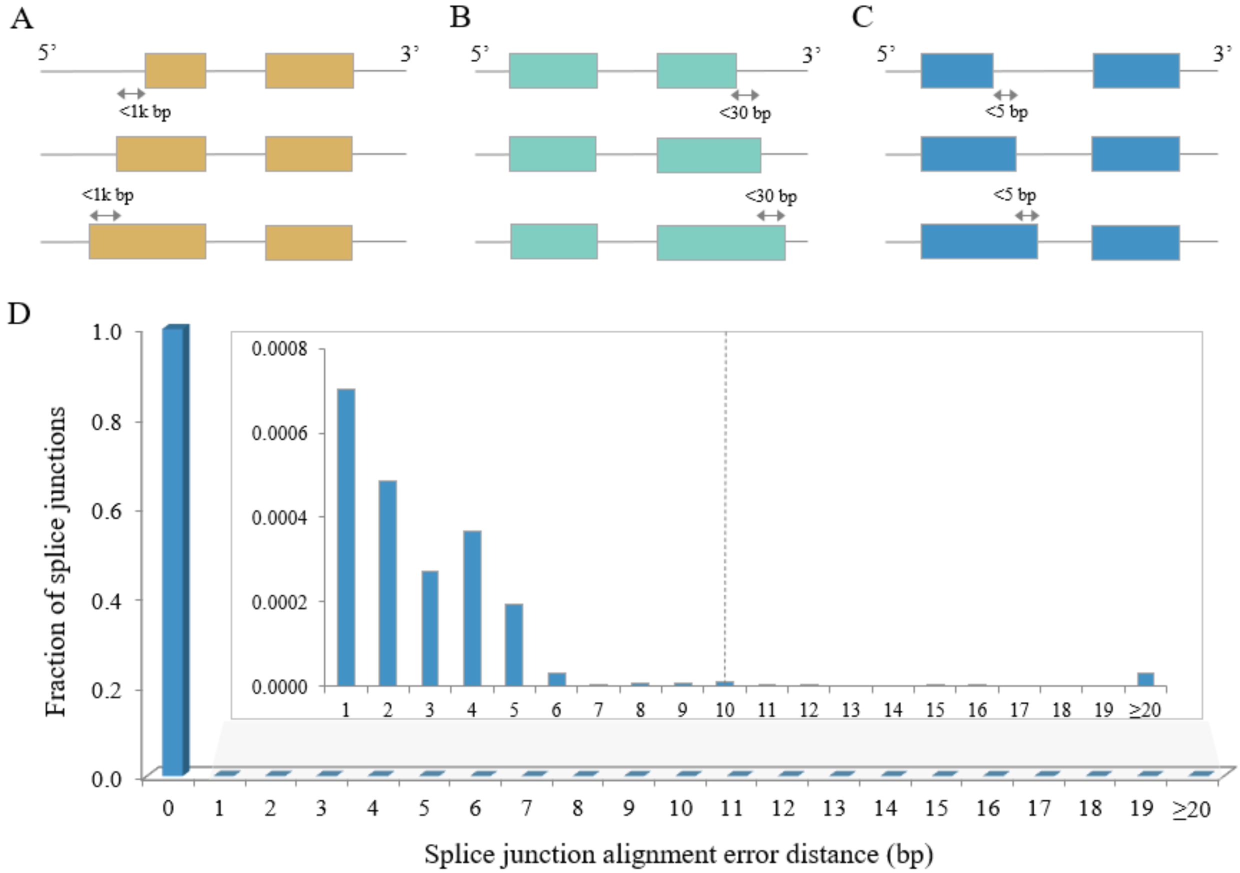


**Figure S3 Schematic diagram for the collapsing of redundant transcript models.** The criteria to collapse similar transcripts at 5' end, 3' end, and splice junctions are shown in (A) - (C). (D) illustrates the fraction of splice junctions with various alignment error distances to the known junction positions. This result was generated by aligning reference annotated cDNA sequences (*i.e.*, with known splice junction positions) back to the reference genome. The dashed line marks the cutoff (10bp) of the alignment error distance of splice junctions to collapse similar transcripts. Given that collapsing process occurs in two directions, the allowed splice junction difference is set as 5bp, as shown in (C).

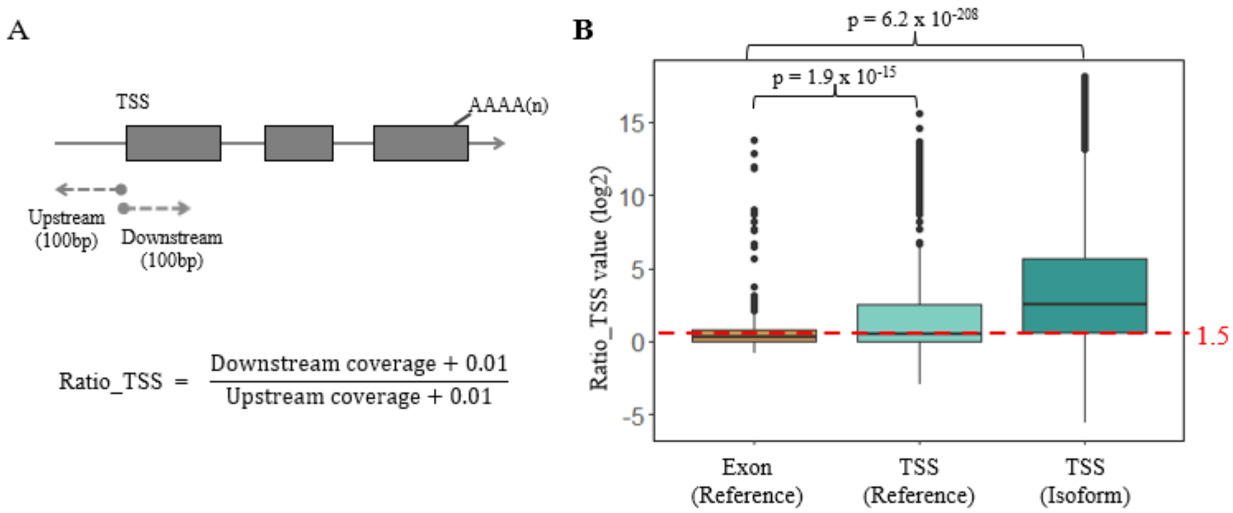


**Figure S4 Calculation of coverage ratios of TSSs.** (A) depicts the methodology to calculate the ratio coverage for a given TSS. The Ratio_TSS is calculated as the ratio (inside coverage + 0.01) / (outside coverage + 0.01), where the mean coverage of 100bp upstream of TSS is taken as coverage inside isoform and that of 100bp downstream as outside coverage (Tardaguila et al. 2018). (B) shows the comparison of Ratio_TSS distributions across different sets of TSSs. Boxes represent the interquartile range (IQR, distance between the first and third quartiles), with black lines in the middle to denote the median. The boundaries of the whiskers are based on the 1.5 IQR values for both sides, and black dots represent outliers. The 1,000 randomly chosen sites from exon regions annotated in GRCm39/mm39 reference based on Ensembl v103 were considered as negative control dataset, and 1,000 randomly chosen TSSs from reference annotation as positive control dataset, and all the TSSs of the filtered PacBio Iso-Seq isoforms detected in our study as the test dataset. For each assigned site, ratio_TSS represents the maximum value of the ratios across all 48 RNA-Seq samples. The red dashed line marks the threshold of Ratio_TSS (median of values from positive control dataset, *i.e.*, 1.5) to define "reliable" TSSs. Statistical p-values were computed using two-sided Wilcoxon rank sum tests.

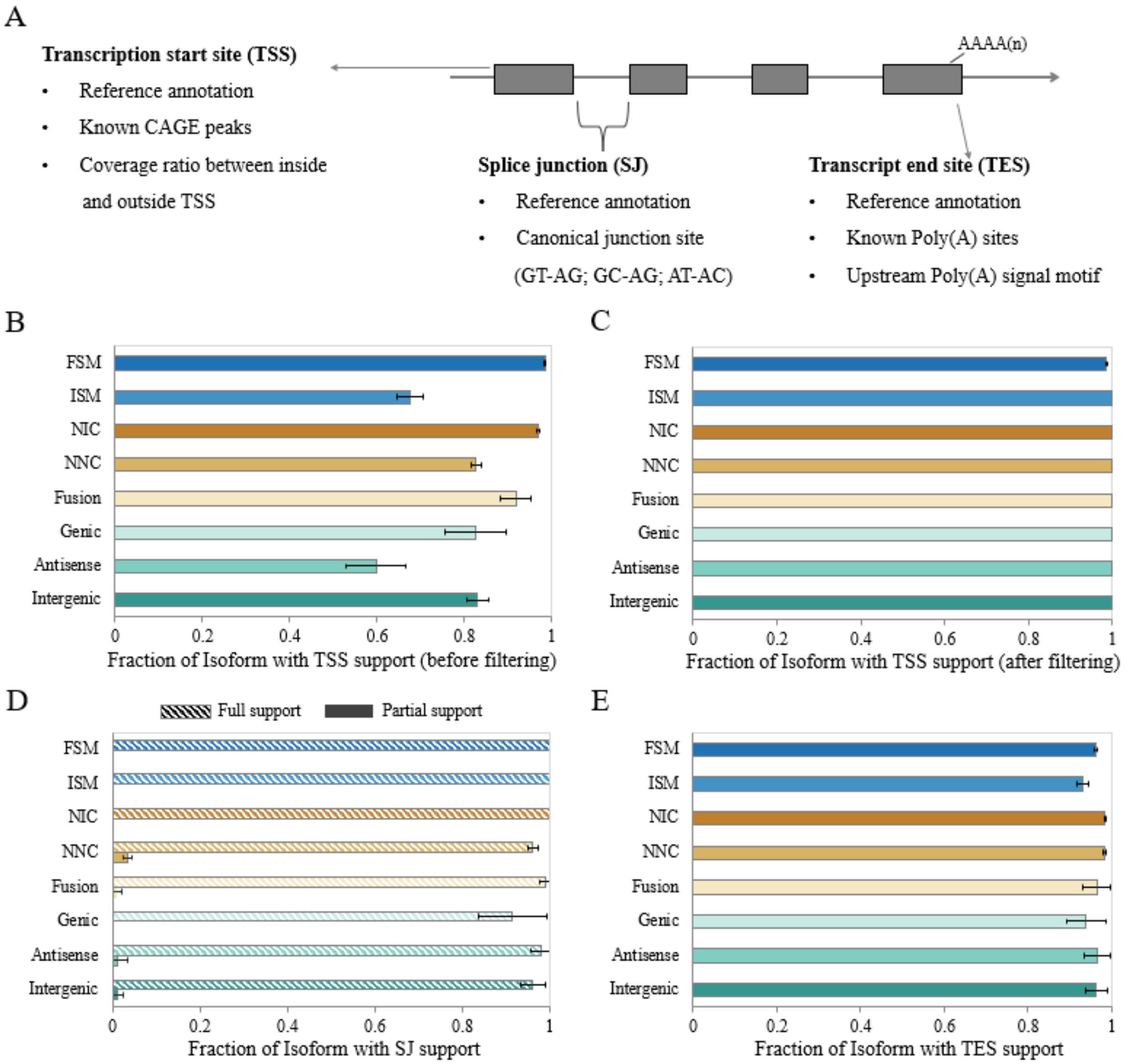


**Figure S5 Support level for transcripts from different structural categories.** (A) depicts the evidence lines to define reliable isoforms concerning TSS, SJ, and TES. (B) and (C) show the TSS support evidence before and after applying TSS filtering criteria on non-FSM isoforms. (D) and (E) show the support evidence for SJ and TES, respectively. Full SJ support indicates that all the SJs within the focal isoform are supported, and partial support for the ones with at least one supported SJ. The error bars indicate the standard error of means (SEM) of the fraction support values.

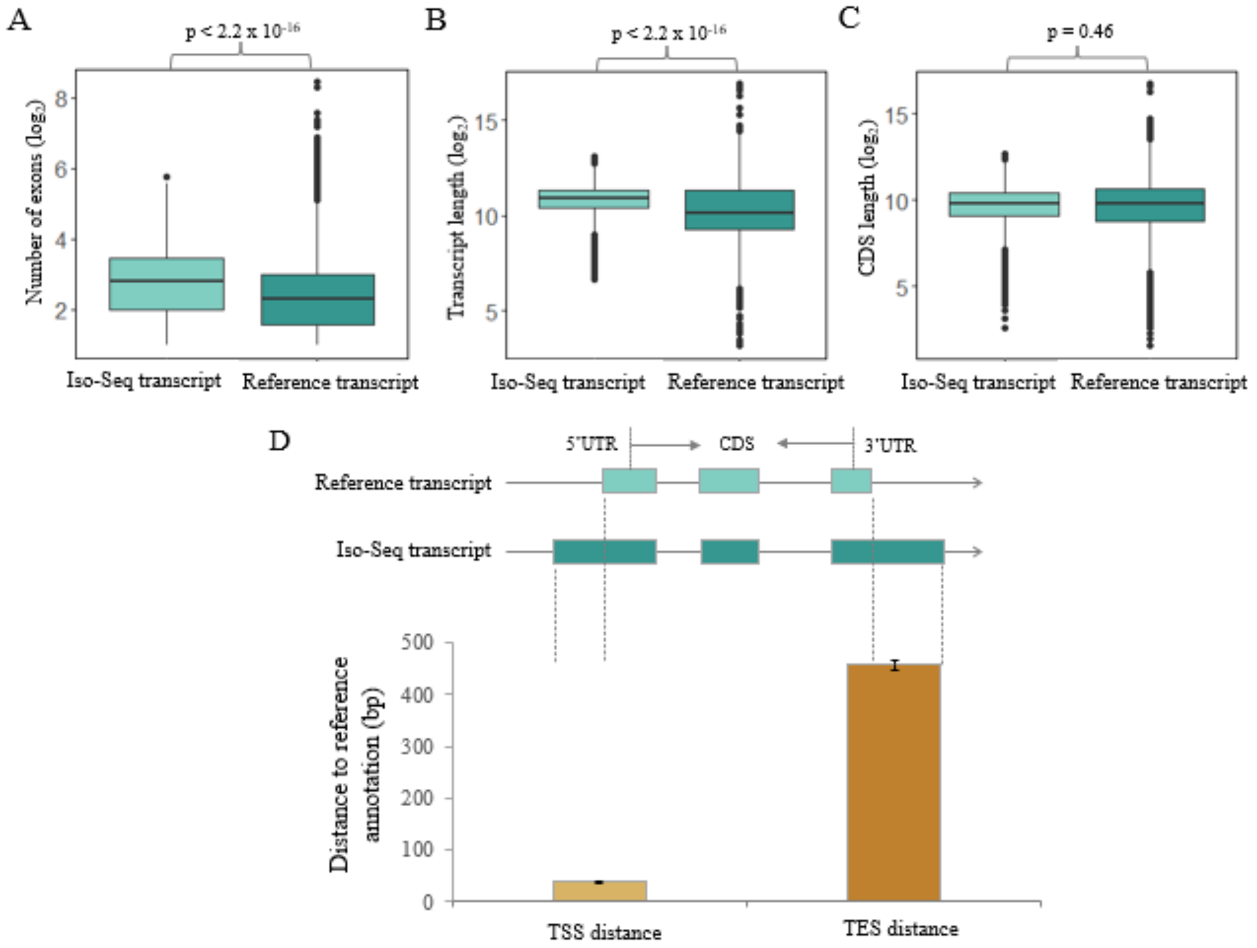


**Figure S6 Features of all Iso-Seq detected transcripts compared to reference annotated transcripts.** (A)–(C) show the frequency distribution of exon number, transcript length, and CDS length between Iso-Seq defined transcripts and reference annotated transcripts. Boxes represent the interquartile range (IQR, distance between the first and third quartiles), with white dots (or black lines) in the middle to denote the median. The boundaries of the whiskers are based on the 1.5 IQR values for both sides, and the black dots represent outliers. Statistical p-values were computed using two-sided Wilcoxon rank sum tests. (D) indicates the distribution of distances to the transcription start sites and termination sites between Iso-Seq detection and reference annotation. Error bars show the features' standard errors of the mean (SEM) values. Only Iso-Seq transcripts matching perfectly to reference annotated transcripts (FSM) were analyzed here.

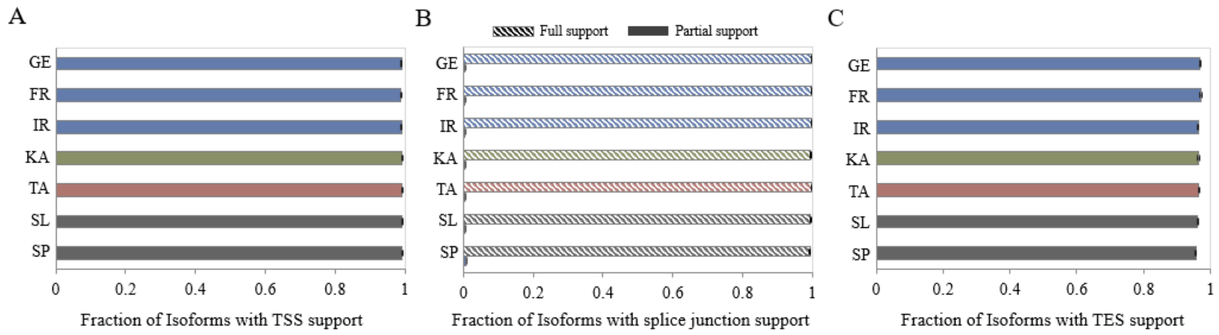


**Figure S7 Support level of transcripts from different populations.** (A), (B) and (C) show the support evidence for TSS, SJ, and TES, respectively. Full SJ support indicates that all the SJs within the focal isoform are supported, and partial support for the ones with at least one supported SJ. The color code for each population follows Figure 2, and the description of supporting evidence for reliable isoforms can be found in Supplemental Figure S5. The error bars indicate the standard error of means (SEM) of the fraction support values.

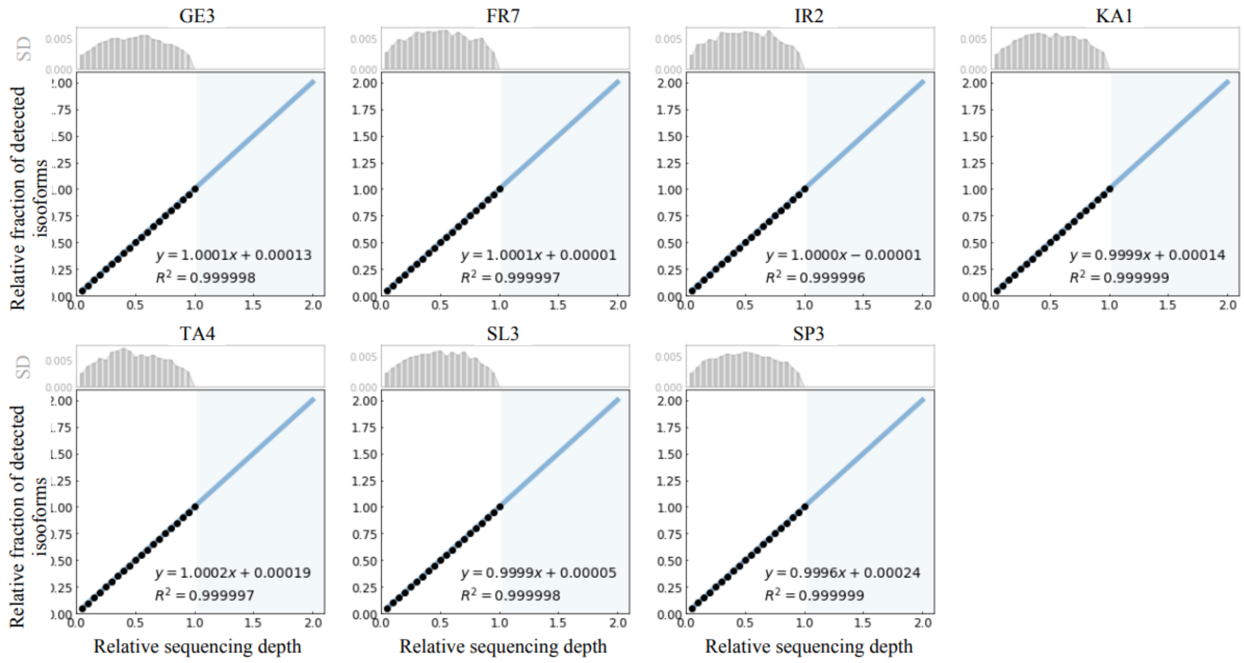


**Figure S8 Rarefaction analysis for singletons from individuals of the different populations**. The singleton isoforms were defined as those supported by singleton FLNC read. The resampling sequencing depth was selected from 0.05 to 1, with a step size 0.05. The blue area shows the prediction after doubling the current Iso-Seq sequencing depth. For each of the seven assayed populations, one individual was randomly selected and analyzed as a representative. The bar plots in the above panels show the standard deviation (SD) of each resampling analysis.

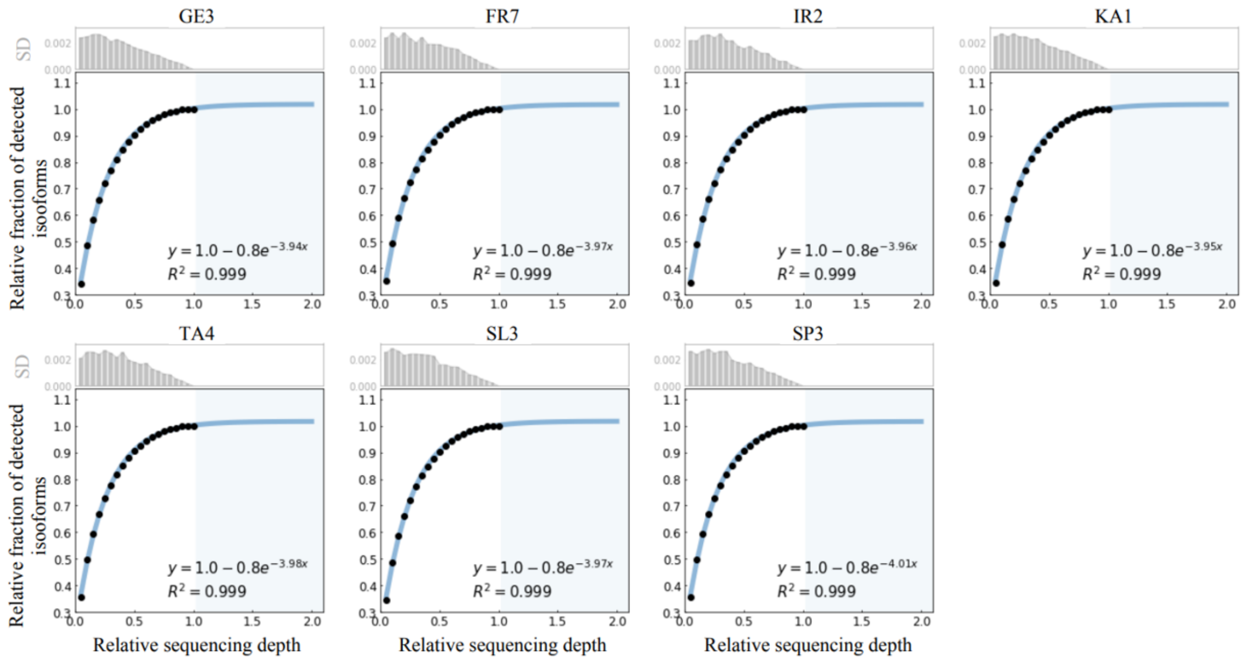


**Figure S9 Rarefaction analysis for high confidence isoforms from individuals of the different populations**. The high-confidence isoforms are defined as those supported by at least two FLNC reads. The resampling sequencing depth was selected from 0.05 to 1, with a step size 0.05. The blue area shows the prediction after doubling the current Iso-Seq sequencing depth. For each of the seven assayed populations, one individual was randomly selected and analyzed as a representative. The bar plots in the above panels show the standard deviation (SD) of each resampling analysis.

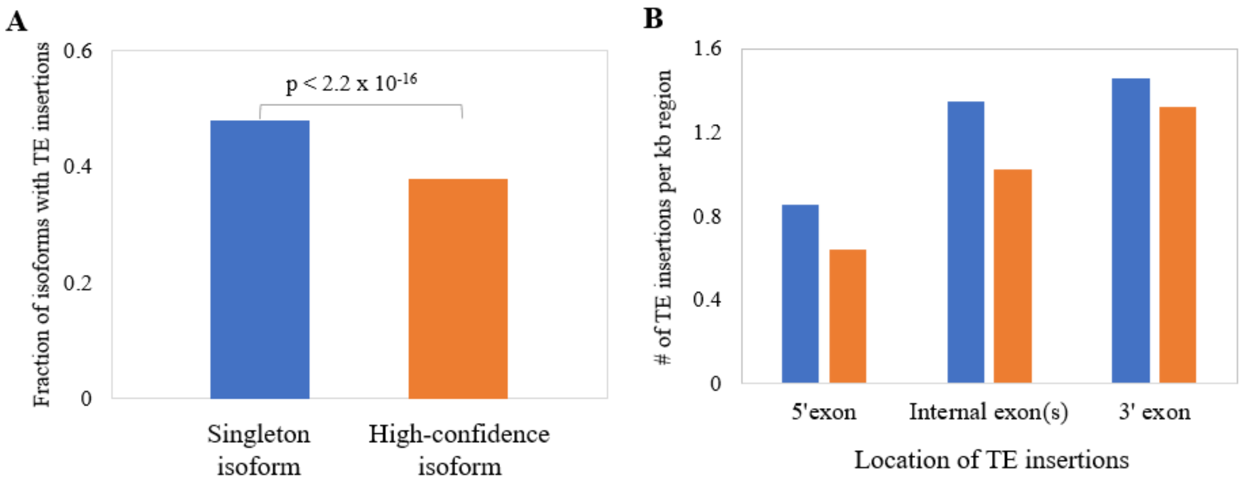


**Figure S10 Comparison of TE insertions between singleton isoforms and high-confidence isoforms.** (A) show the fractions of isoforms with at least one TE insertion for singleton and high-confidence isoforms. The statistical test was performed using Fisher's exact test. (B) show the distributions of TE insertion locations for singleton and high-confidence isoforms. Only isoforms with at least one internal exon (i.e., at least 3 exons in total) were analyzed. The number of TE insertions for each exon was normalized on the basis of the exon length.

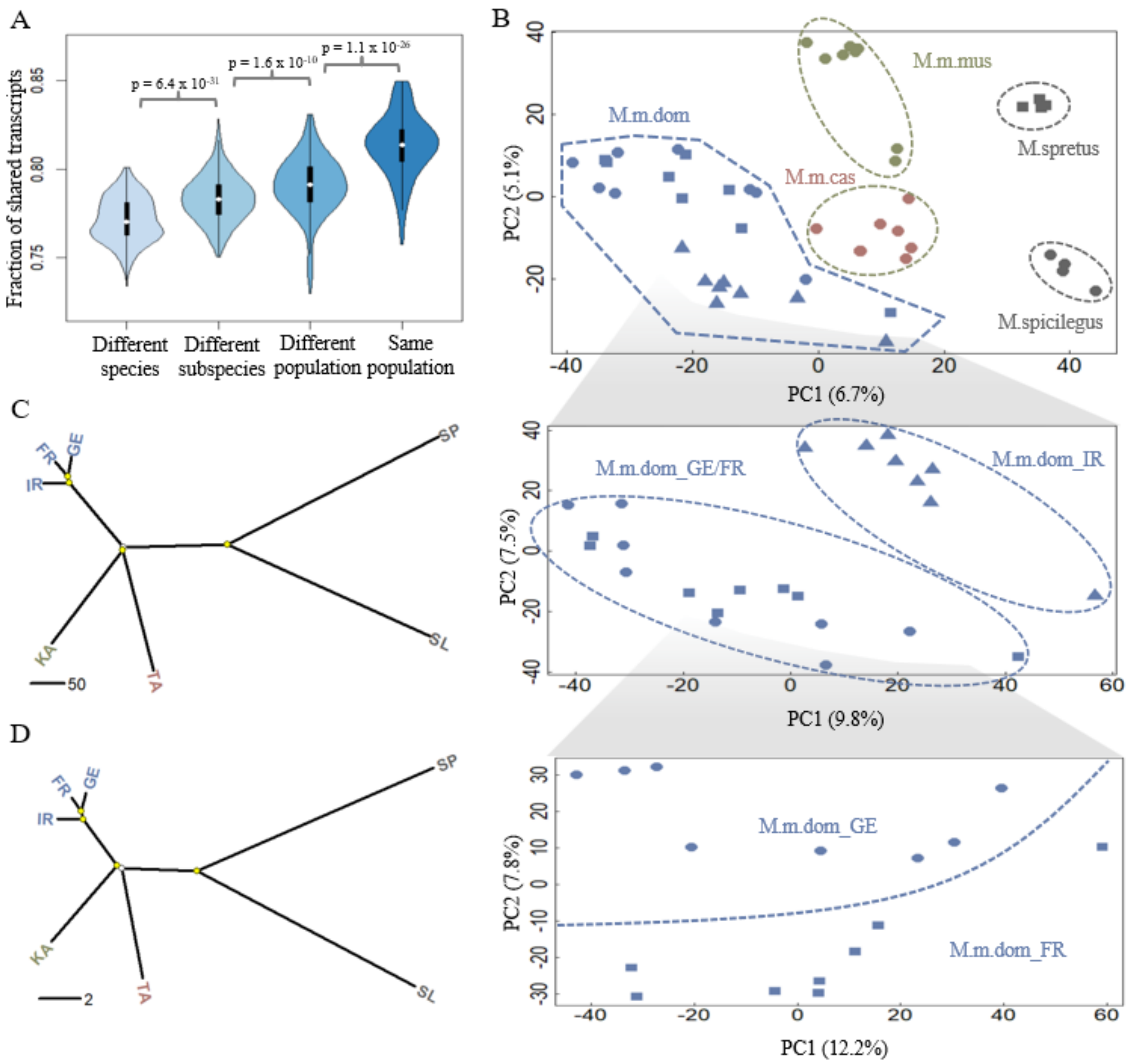


**Figure S11 Population differentiation pattern based on higher-frequency isoforms.** This figure is related to Figure 2, but only isoforms present in at least two mice individuals were included for the analysis. (A) shows the fraction of shared isoforms for pair-wise individual comparison. The fraction of shared isoforms between each pair of individuals is defined as the number of overlapping isoforms divided by the average number of detected isoforms of the two individuals to compare. Black boxes represent the interquartile range (IQR, distance between the first and third quartiles), with white dots in the middle to denote the median. The boundaries of the whiskers (also the ranges of violins) are based on the 1.5 IQR values for both sides. The statistical p-values of the fractions of shared isoforms between different comparison groups were computed using Wilcoxon rank sum tests. (B) shows the projection of the top two PCs of isoform variation in house mouse and outgroup individuals. Enlarged insets represent the results for the three populations from the subspecies of *M. m. domesticus*, as they cannot be well distinguished in the main figure. (C) and (D) show phylogenetic trees built on the basis of isoform and SNP variants fixed within each population, respectively. Split nodes marked in yellow are the ones with bootstrap support value >70%.

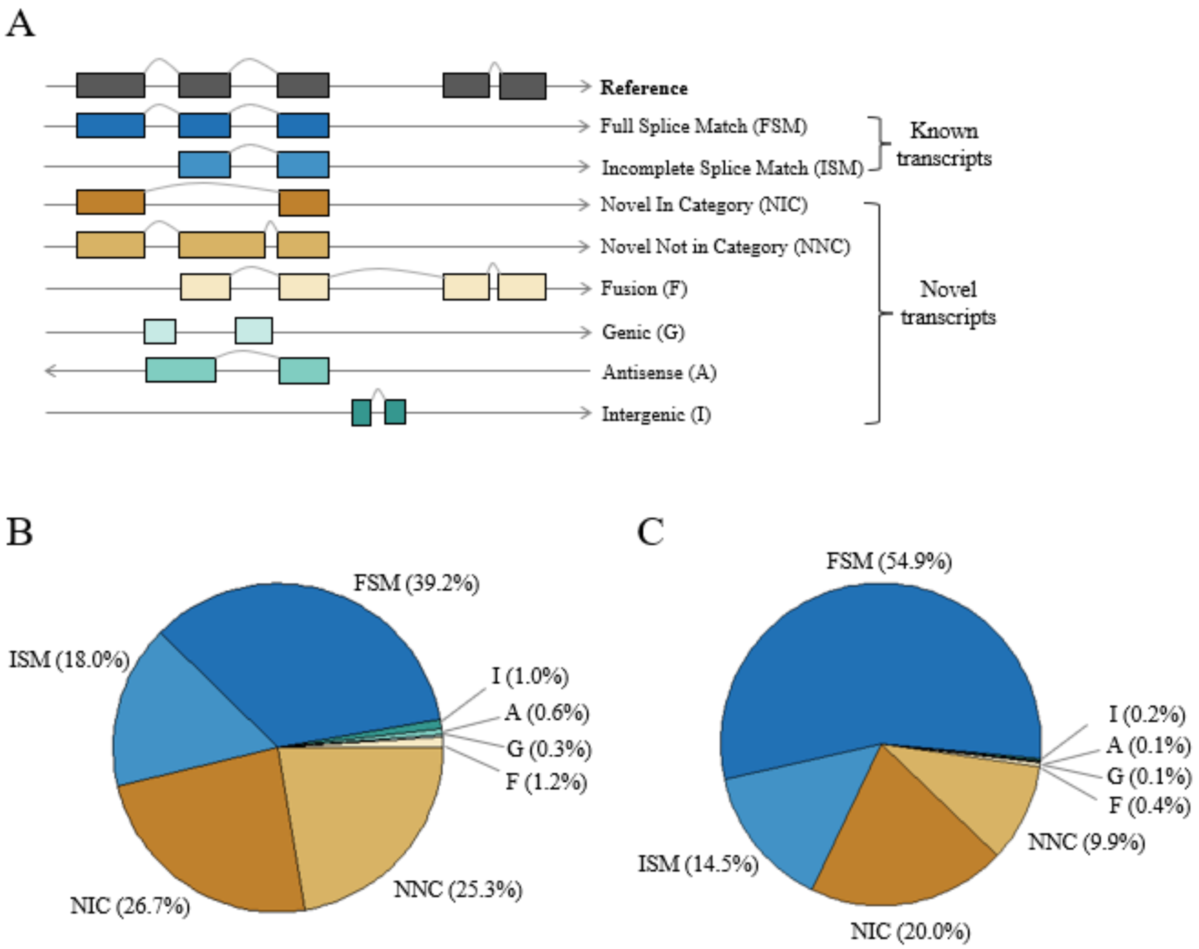


**Figure S12 Distribution of structural categories of house mouse isoforms.** (A) Types and illustrations of the isoforms. (B) Fraction distribution of structural categories of isoforms detected in house mouse populations (*i.e.*, excluding the outgroup specific isoforms). (C) Fraction distribution of structural categories of isoforms conserved in house mouse and outgroup species.

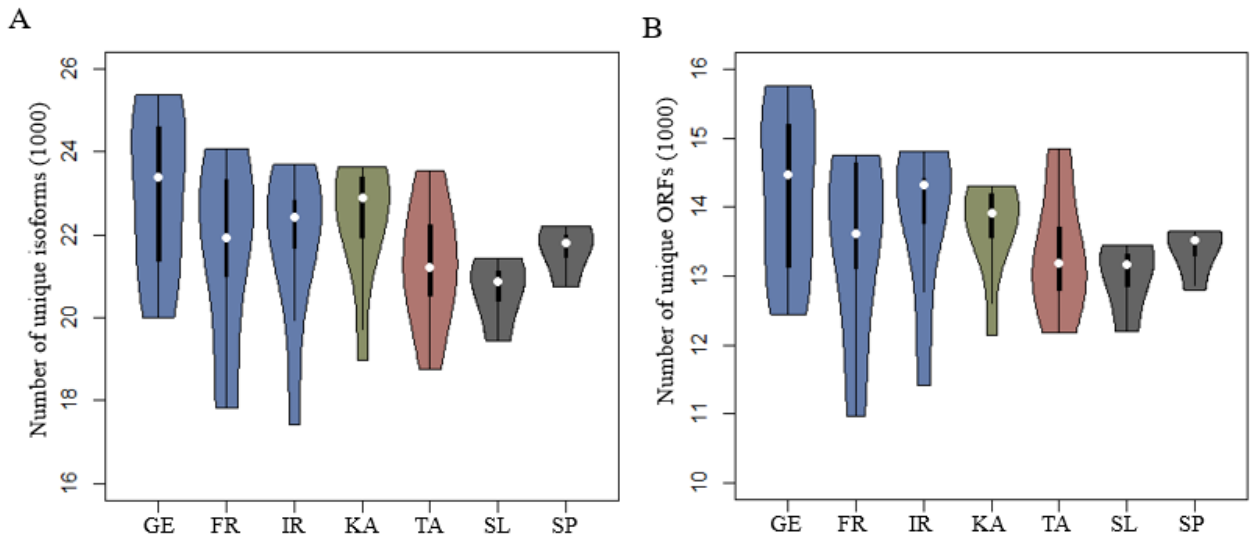


**Figure S13 Distribution of isoform and ORF numbers across all the seven assayed populations.** (A) shows the distribution of the numbers of unique isoforms, and (B) shows the distribution of the number of unique ORFs. Black boxes represent the interquartile range (IQR, distance between the first and third quartiles), with white dots in the middle to denote the median. The boundaries of the whiskers (also the ranges of violins) are based on the 1.5 IQR values for both sides.

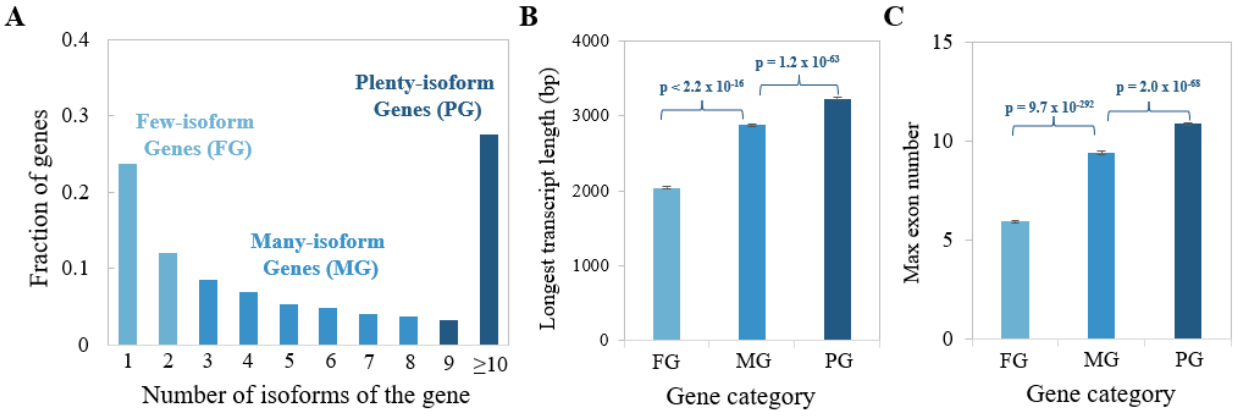


**Figure S14 Genomic features of gene loci with different numbers of isoforms.** (A) Gene categories were based on the number of isoforms of each gene. Three gene categories were classified with roughly the same number of genes in each group. (B) Comparison of the length of the longest transcript for a given gene among the three gene categories. (C) Comparison of the maximum number of exons across all the detected transcripts for a given gene among the three gene categories. Error bars show standard errors of the mean (SEM) values. Statistical p-values were computed using two-sided Wilcoxon rank sum tests.

**Table S1 Summary of sequencing statistics generated in this study.** Details on the sequencing statistics for each individual can be found in Supplemental Data S1B and S1C.

| Species | Subspecies | Population | # of sampled individuals | Sex | Organ | PacBio Iso-Seq # of raw sub-reads ($10^6$) | Illumina RNA-Seq # of raw read pairs ($10^6$) |
|---|---|---|---|---|---|---|---|
| Mus musculus | M. m. domesticus | Germany (GE) | 8 | Male | Whole brain | 55.8 (SD:8.1) | 27.1 (SD:3.0) |
| | | France (FR) | 8 | Male | Whole brain | 54.2 (SD:13.2) | 26.7 (SD:2.5) |
| | | Iran (IR) | 8 | Male | Whole brain | 61.5 (SD:13.2) | 26.3 (SD:1.1) |
| | M. m. musculus | Kazakhstan (KA) | 8 | Male | Whole brain | 61.0 (SD:8.5) | 29.6 (SD:2.1) |
| | M. m. castaneus | Taiwan (TA) | 8 | Male | Whole brain | 59.8 (SD:7.4) | 29.6 (SD:5.2) |
| Mus spicilegus | M. spicilegus | Slovakia (SL) | 4 | Male | Whole brain | 69.1 (SD:6.5) | 30.4 (SD:2.0) |
| Mus spretus | M. spretus | Spain (SP) | 4 | Male | Whole brain | 58.1 (SD:3.4) | 29.7 (SD:4.4) |

**Table S2 Summary of statistics for the comparison analysis between known and novel transcripts.** Known transcripts are the ones from the categories "FSM" and "ISM", and Novel transcripts from the categories "NIC", "NNC", "F", "G", "A", and "I". Exon-derived novel transcripts are the ones with categories "NIC", "NNC", and "F"; others are defined as ones with categories "G", "A", and "I". Fisher's exact tests were exploited for fraction comparison analyses, and two-sided Wilcoxon rank sum tests for the rest analyses. NMD: nonsense-mediated decay. NMD analysis was performed on coding transcripts only.

| | **Known vs. Novel** | | **Exon-derived Novel vs. Others** | |
|---|---|---|---|---|
| | **Mean value** | **p-value** | **Mean value** | **p-value** |
| **Exon Number** | 7.6 vs. 8.6 | $< 2.2 \times 10^{-16}$ | 8.8 vs. 2.9 | $< 2.2 \times 10^{-16}$ |
| **Transcript length (bp)** | 2,019 vs. 2,040 | $< 2.2 \times 10^{-16}$ | 2,058 vs. 1,516 | $< 2.2 \times 10^{-16}$ |
| **CDS length (bp)** | 1,056 vs. 976 | $< 2.2 \times 10^{-16}$ | 981 vs. 538 | $< 2.2 \times 10^{-16}$ |
| **Fraction of coding transcripts** | 0.90 vs. 0.87 | $3.0 \times 10^{-78}$ | 0.89 vs. 0.32 | $< 2.2 \times 10^{-16}$ |
| **Fraction of NMD transcripts** | 0.02 vs. 0.18 | $< 2.2 \times 10^{-16}$ | 0.20 vs. 0.15 | $4.2 \times 10^{-4}$ |
| **# of individuals with expression (TPM > 0)** | 13.01 vs. 4.66 | $< 2.2 \times 10^{-16}$ | 4.73 vs. 2.78 | $5.7 \times 10^{-37}$ |
| **Expression level (TPM)** | 15.66 vs. 1.05 | $< 2.2 \times 10^{-16}$ | 1.06 vs 0.66 | $1.9 \times 10^{-31}$ |

### SI References


Abugessaisa I, Noguchi S, Hasegawa A, Kondo A, Kawaji H, Carninci P, Kasukawa T. 2019. refTSS: A Reference Data Set for Human and Mouse Transcription Start Sites. *J Mol Biol* **431**: 2407-2422.

Derti A, Garrett-Engele P, Macisaac KD, Stevens RC, Sriram S, Chen R, Rohl CA, Johnson JM, Babak T. 2012. A quantitative atlas of polyadenylation in five mammals. *Genome Res* **22**: 1173-1183.

Herrmann CJ, Schmidt R, Kanitz A, Artimo P, Gruber AJ, Zavolan M. 2020. PolyASite 2.0: a consolidated atlas of polyadenylation sites from 3' end sequencing. *Nucleic Acids Res* **48**: D174-D179.

Howe KL, Achuthan P, Allen J, Allen J, Alvarez-Jarreta J, Amode MR, Armean IM, Azov AG, Bennett R, Bhai J et al. 2021. Ensembl 2021. *Nucleic Acids Res* **49**: D884-D891.

Kuo RI, Cheng Y, Zhang R, Brown JWS, Smith J, Archibald AL, Burt DW. 2020. Illuminating the dark side of the human transcriptome with long read transcript sequencing. *BMC Genomics* **21**: 751.

Leung SK, Jeffries AR, Castanho I, Jordan BT, Moore K, Davies JP, Dempster EL, Bray NJ, O'Neill P, Tseng E et al. 2021. Full-length transcript sequencing of human and mouse cerebral cortex identifies widespread isoform diversity and alternative splicing. *Cell Rep* **37**: 110022.

Tardaguila M, de la Fuente L, Marti C, Pereira C, Pardo-Palacios FJ, Del Risco H, Ferrell M, Mellado M, Macchietto M, Verheggen K et al. 2018. SQANTI: extensive characterization of long-read transcript sequences for quality control in full-length transcriptome identification and quantification. *Genome Res* doi:10.1101/gr.222976.117.

Xu C, Park JK, Zhang J. 2019. Evidence that alternative transcriptional initiation is largely nonadaptive. *PLoS Biol* **17**: e3000197.

Xu C, Zhang J. 2018. Alternative Polyadenylation of Mammalian Transcripts Is Generally Deleterious, Not Adaptive. *Cell Syst* **6**: 734-742 e734.
